# Contact heterogeneity in network reconstruction: evaluating synthetic graph generators for rabies transmission modeling

**DOI:** 10.64898/2026.10.16.759544

**Authors:** Inez Derkx, James Mac Hyman, Fatima Abdelrazak Zakaria, Mahamat Fay Abakar, Salome Dürr, Nakul Chitnis

## Abstract

Canine rabies remains endemic across much of Africa and Asia, causing approxi 59,000 global deaths yearly. Network-based models that reconstruct contact pat are central to predicting canine rabies transmission and to designing targeted interventions, but the choice of network generator algorithm may alter network structure and outbreak predictions. We therefore compared five network generators using five empirical free-roaming dog contact networks from Chad, Indonesia, an Guatemala in network topology and outbreak dynamics under susceptible-exposed-infectious-removed (SEIR) and susceptible-infectious-suscep (SIS) transmission across a range of infectious-period lengths.

We found that the degree-corrected stochastic block model (DCSBM) most consistently reproduces both empirical topology and outbreak dynamics, achievi Mahalanobis distance of 17.6 to the empirical Chad network compared to a rang 21.0 to 220.7 across the other generators and yielding similar results for the othe locations. Moreover, the SIS and SEIR outbreak dynamics reveal that the DCSB overall best reproduces the simulated empirical outbreak dynamics, but that the consequences of generator choice are not uniform: network reconstruction matter precisely for short-infectious-window pathogens like canine rabies and progressive for long-infectious-window pathogens. By an infectious window of nine days, the outbreak dynamics across four of five generators converge to within two percent points of the empirical reference, even when their underlying topologies differ substantially. This pathogen-dependent sensitivity offers a principled criterion for generator selection is consequential and when it is not.

This work reiterates the importance of both degree heterogeneity and commu structure for faithful contact-network reconstruction in canine rabies modeling, particularly for short-window infectious diseases where contact heterogeneity drives transmission. For longer-window pathogens, simpler generators are likely adequate.

**Author summary:** Canine rabies causes around 59,000 deaths worldwide each year, and its spread l depends on the contact networks domestic dogs form with each other. In these networks, some dogs have far more contacts than others, and this heterogeneous structure shapes how diseases like rabies are spread through a population. These structures can be reconstructed using different models, though the choice of model impacts how well it can reconstruct an observed contact network. Here, we ther compare five synthetic network models against real contact data collected from Chad, Indonesia, and Guatemala. We evaluate both the structure of the network generate and how well they reproduce outbreak dynamics, comparing the outbreaks resulting from the networks generated by each model to outbreaks simulated on observed networks themselves. One model, the degree-corrected stochastic block best reproduces the observed contact structures across locations and comes close matching outbreak simulations based on the observed network. We also find that choice matters more when pathogens have short infectious periods: the models d more in their outbreak outcomes under short infectious windows and all but one b more similar when infectious periods are longer.

## Introduction

Contact patterns play a vital role in the spread of infectious diseases [1–3]. While compartmental population models of disease spread traditionally assume homogeneity in contact patterns [4], population contact structures are generally heterogeneous and impact the duration, size and dynamics of epidemic outbreaks [5–10].

Incorporating heterogeneous contact structures in disease models has highlighted how the social positions of individuals and their clustered interactions can alter disease transmission patterns and help in designing targeted intervention strategies [3, 11–14]. For instance, individuals with a greater number of contacts (degree centrality) tend to be disproportionately exposed to pathogens and therefore pose a high risk to epidemic outbreaks; and degree heterogeneity can substantially lower or even remove the conventional epidemic threshold of mean-field epidemiological models [13, 14]. In networks with strong community structure, targeting individuals who function as bridges between communities (betweenness centrality) has been shown to be an effective intervention strategy [15–18]. In addition to individual-scale heterogeneity, network clustering (global clustering coefficient) can also profoundly impact epidemic dynamics, with increased clustering leading to more local spread but simultaneously a reduction in epidemic growth rate [19–21].

Although capturing the heterogeneity of population contact structures is therefore required to model disease dynamics, this effort also relies on the accuracy of empirical proximity-based networks and their ability to represent pathogen transmission opportunities. This requirement is best met for pathogens transmitted exclusively via direct contact or close proximity, such as rabies [22, 23]. Canine rabies is a zoonotic disease responsible for approximately 59,000 yearly human deaths globally and is still endemic in many countries in Africa and Asia [24, 25]. Different network-based approaches have successfully used contact structures to model canine rabies transmission, for instance linking its persistence to social disruptions caused by rabies-induced behavioral changes [26] and providing support for network centrality-based targeted vaccinations in domestic dog populations [3].

In the latter study, Laager et al. [3] generated synthetic population networks from domestic dog proximity data using a spatially explicit network construction algorithm, with model outcomes matching city-wide rabies outbreak data from N’Djamena, Chad. Even though empirical degree distribution guided edge formation, other sources of heterogeneity were not considered. Given the impact of degree and community structure on epidemic dynamics, it is likely crucial that network generator algorithms encapsulate not only local (individual-level), but also global (network-level) contact variation [27, 28]. However, it remains unclear whether incorporating different or additional sources of heterogeneity in network reconstruction improves rabies transmission prediction.

Numerous generative models exist that could address this gap and have been used to reconstruct empirical networks, and which can optimize both local and global network characteristics [29–33]. These include random graph generators without population structure (e.g., Erdős–Rènyi graphs [34]), scale-free network models (e.g., Barabási–Albert model [35]), models optimized to produce specific local or global structural properties such as degree distribution (e.g., Chung–Lu model [36]), community structure (e.g., stochastic block model and variations thereof [37]), or clustering through triadic closure (e.g., clustered random-graph model [38, 39]). The successful application of these models highlights their potential as synthetic network generators for disease modeling. Yet, systematic comparisons of how different models, including the previously used model of Laager et al. [3], capture empirical contact heterogeneity and the consequences thereof for rabies modeling are still lacking.

This paper therefore assesses the role of contact heterogeneity in network reconstruction for rabies modeling and asks a more general question: when does the choice of network generator actually matter for outbreak prediction, and when does it not? We compare five network generator algorithms against an empirical contact network of dogs residing in N’Djamena, Chad, and across four additional locations in Indonesia and Guatemala. Specifically, we examine a) how the synthetic networks compare in network topology to their empirical references, b) how the variation in graph heterogeneity impacts disease transmission in outbreak simulations of an SEIR rabies model and an SIS model (e.g., common cold) for comparison, and c) whether these findings generalize when applied to other variably sized empirical contact networks. This work extends the rabies-modeling study of Laager et al. [3], which evaluated a single spatially explicit network construction algorithm (SENCA), by placing four additional generator families alongside it and testing whether their alternative encoding of heterogeneity improves transmission-prediction fidelity. It also builds on recent comparative work on stochastic block models for empirical contact networks [32] by pairing topology comparison with full SEIR and SIS outbreak simulations across multiple infectious-period regimes, rather than comparing topology alone.

While numerous alternative models could be explored for this purpose, we limit our comparison to five models which in combination examine the relative importance of three axes of structural heterogeneity: degree-distribution, community structure, and explicit triadic-closure (clustering) targeting, allowing us to assess the incremental contribution of each structural feature to network reconstruction and simulated transmission. As previously stated, these three axes have distinct epidemiological implications, including individual contribution to acquisition and transmission of infection (degree), extent of localization versus universal spreading (community structure), and capturing contact redundancy along with determining infection propagation speed (clustering).

The models therefore include the SENCA, as introduced above, which is optimized to fit an empirical degree distribution [3], a stochastic block model (SBM) which aims to reconstruct empirical community structure [37], a degree-corrected stochastic block model (DCSBM) which fits both degree heterogeneity and community structure [40], a Newman clustered random-graph generator (NCRG) which additionally targets empirical levels of triadic closure [39], and an Erdős–Rènyi model (ERM) with random contact structure [34] as a control. For the remainder of the article, we also refer to these five network generators as ‘graph types’.

## Materials and methods

### Free-roaming dog proximity networks

We built an empirical contact network from interactions between free-roaming domestic dogs in an urban area covering approximately 1 km^2^ in N’Djamena, Chad. These data were collected in 2016 using tailor-made geo-located contact sensors attached to dog collars over the span of approximately 3.5 days. Rather than attempting to sample a single fully connected network, as many dogs as possible inhabiting the covered study area were sampled. The resulting network may therefore include isolated individuals or sub-groups. While GPS coordinates of the home locations of all dogs as well as individual GPS locations were also collected, these were not included in contact or network formation. We used data collected in district six of the city to create an undirected, unweighted simple contact network of 235 dogs. In this network, two nodes (dogs) are connected by an edge (contact) if they were observed in close proximity at least once during the study period. This network is referred to as ‘empirical network’ throughout the remainder of the manuscript. Further information about the data collection can be found in detail in Laager et al. [3].

In addition to this empirical network, we examined previously published contact data of free-roaming domestic dogs in Indonesia and Guatemala collected in 2018 for further comparison [41]. The study areas include two rural sites in Guatemala: La Romana and Sabaneta; and one rural and one semi-urban site in Indonesia: Hepang and Habi, respectively. Data collection followed a similar protocol compared to the 2016 study in Chad, including how the area of the study sites was selected, how the dogs were sampled, and how contacts were established using geo-located sensors. Further detailed descriptions of the study sites and data collection can be found in Warembourg et al. [41]. While these networks are also empirical contact networks, we refer to them throughout the manuscript by their location names instead (e.g., the Sabaneta network).

### Network generator algorithms

We generated networks using five models representing distinct combinations of degree-distribution heterogeneity, community structure, and clustering (Table 1). Detailed specifications of each generator are provided below.

**Table 1.** Comparison of the five graph generators and their structural properties. Columns give the network type, how links between nodes are created, and whether the degree distribution, community structure, and explicit triadic-closure (clustering) targeting are modeled. The fifth, sixth and seventh columns summarize each generator’s structural signature.

| Graph type | Abbrev. | Network type | Edge creation | Degree dist. | Community structure | Cluster. target | Source |
| --- | --- | --- | --- | --- | --- | --- | --- |
| Spatial | SENCA | Assortative mixing | Spatial and preferential attachment of links | Heterog. | No | No | Laager et al. [3] |
| Stochastic Block | SBM | Assortative mixing | Based on community membership | Homog. (within comm.) | Yes | No | Holland et al. [37] |
| Degree-Corrected Stochastic Block | DCSBM | Assortative mixing | Community membership and node degree | Heterog. | Yes | No | Karrer & Newman [40] |
| Newman Clustered Random Graph | NCRG | Triangle-augmented configuration | Single-edge and triangle-edge stubs matched to empirical degree and clustering | Heterog. | No (implicit) | Yes (explicit) | Newman [39] |
| Erdős-Rényi | ERM | Random | Random | Homog. | No | No | Erdős & Rényi [34] |

First, we tested a previously published spatially explicit network construction algorithm (SENCA), which was designed to model rabies outbreak dynamics by generating a similar degree distribution to the empirical network it is based on [3]. In short, this algorithm requires a pre-defined number of nodes, assigns them coordinates using random sampling, and implements a two-step process starting with spatial edge construction followed by preferential edge attachment optimized to minimize the Kolmogorov-Smirnov distance between the generated and empirical degree distributions. This algorithm has been successfully used to model rabies outbreaks, with results matching real-world outbreak dynamics and vaccination coverage in N’Djamena, Chad. A more detailed breakdown of the algorithm and its previous implementations can be found in Laager et al. [3].

Furthermore, we explored the performance of a stochastic block model (SBM) [37] and one of its extensions: the degree-corrected stochastic block model (DCSBM) [40]. The SBM is built on the premise that the probability of an edge between two nodes relies solely on the communities (‘blocks’) to which those nodes belong and hence assumes homogeneity between nodes within the same communities. In empirical networks, these communities could for instance reflect geographical neighborhood clusters or dog packs that more frequently interact, though they are treated as latent properties of the network in this study. The DCSBM introduces a degree parameter *ψ* that represents the propensity of nodes to form edges independent of the community to which they belong. It shows the ratio of node degree to its group’s expected degree, hence removing within-community node homogeneity [42]. This allows for examination of node-level variability in contacts, for instance because some dogs roam further than others and therefore have more contacts, which was highlighted as an important factor in rabies transmission in previous work [3]. While the SBM thus introduces heterogeneity through community structure, the DCSBM additionally models node degree.

For both block models, the optimal number of communities *K* is estimated from the empirical adjacency matrix *A* by asymptotic likelihood ratio [43]. Nodes are then allocated to one of *K* communities through regularized spherical spectral partitioning (RSSP). RSSP is a community detection algorithm that applies regularization to either the adjacency matrix (if SBM) or the Laplacian matrix (if DCSBM) by adding a constant to each element in these matrices, after which clustering is performed on the spherical coordinates that are assigned to each node [44, 45]. The resulting community (block) labels *g* and the estimated block connection probability matrix *B* are subsequently used to generate the synthetic graph for the SBM, where the probability *P_ij_* of an edge between nodes *i* and *j*is *B_gigj_* . The DCSBM instead uses the estimated probability matrix *P*^^^, calculated using *B*, *g*, and *ψ*, to generate a graph in which *P_ij_* = *ψ_i_ψ_j_B_gigj_* . The process of selecting and assigning *K* as well as estimating *B*, *ψ* and *P*^^^ is performed using the R package *randnet* (v1.0) and all graphs are generated using *igraph* (v2.2.2).

We additionally implemented a Newman clustered random-graph generator (NCRG) following Newman [39]. This generator extends a configuration model, which preserves the empirical degree sequence while otherwise randomizing all connections, to incorporate triangle structure. The NCRG therefore explicitly targets triadic-closure, a feature which none of the other generators provide, while otherwise treating community structure as an emergent consequence of the rewired topology rather than a tunable input. Graph construction relies on a joint degree sequence in which each node *i* is assigned two distinct stub (half-edge) counts: *s_i_* independent-edge stubs and *t_i_* triangle stubs, where *t_i_* reflects the number of triangles node *i* participates in within the empirical network. The joint degree sequence is extracted from a network by counting per-node triangle participations to obtain *t_i_*, with independent-edge stubs computed as *s_i_* = *k_i_ −* 2*t_i_*, where *k_i_* is the total degree of node *i*. Here, the factor of two reflects that each triangle contributes two edges to a node’s degree. Triangle stubs are then grouped into triplets by random shuffling and matched to form triangles, while independent-edge stubs are paired by random perfect matching, following the procedure described by Newman [39]. Because edges can participate in multiple triangles simultaneously, the NCRG’s assumption of edge-disjoint triangles may not always hold. In such cases, any resulting negative *s_i_* values are automatically clamped to zero. The resulting generated graph is simplified by removing self-loops and multi-edges. We translated the *random cluster graph* function of the *NetworkX* Python package (v3.6.1) to R for implementation of this model.

Lastly, we used an Erdős–Rènyi graph model (ERM) as a baseline to test how the empirical network and synthetic heterogeneous networks compare to a random network structure [34]. The R package *igraph* (v2.2.2) was used to implement this model by randomly drawing a predefined number of edges between a predefined number of nodes, which were both set equal to the number of empirical nodes and edges. As a result, the average degree and graph density are identical to the empirical network’s degree and density.

### Network construction and preprocessing

Given that each of the five network generator algorithms is stochastic, we generated multiple graphs per graph type. Each algorithm was run 100, 1000, 2000 and 5000 times with different seeds. Five replicates per seed were generated and confirmed to be identical, after which we selected one graph per seed for analyses. For each resulting graph, we calculated six measures: mean, median, and maximum degree, mean betweenness, global clustering (transitivity) and network density. These were used to examine how the different graph types and network parameters respond to increasing sampling effort (S1 Fig). The empirical graph was included for comparison, but remained consistent across seeds by definition. We employed the Mahalanobis distance metric to examine which graph type showed most similarity to the empirical network across all six measures.

From these six measures, we selected mean degree and mean betweenness to formally examine stochastic convergence across increasingly larger sampling sizes using a running mean with 95% confidence intervals (S2 Fig). The running mean captures how the cumulative (moving) average of these two parameters changes as the number of seeds increases [46]. Convergence was further assessed via the standard error, the stepwise mean drift (the change in mean between successive sample sizes, e.g. 100 to 1000), and the precision gain obtained at each step, reflecting the reduction in the standard error (S1 Table). Based on these convergence metrics, all 5000 graphs per graph type were used for downstream analyses. Each of the five network generators — DCSBM, ERM, NCRG, SBM, SENCA — was therefore run 5000 times, resulting in an equal number of stochastic networks. The resulting distributions of mean betweenness, global clustering, network density, and average, median, and maximum degree were subsequently compared to the empirical network.

Rather than running disease models on a single graph or all 5000 graphs per graph generator, we selected 100 graphs per graph type that best represented the average degree distribution to prevent stochastic graph attributes from skewing modeling outcomes. To do so, we first compared all complete degree distributions of each graph type and computed the Kolmogorov-Smirnov distance to the empirical degree distribution (S3 Fig). We then summarized each graph type’s degree distribution by six statistics computed across all 5000 graphs of that type: the minimum, 25th percentile, mean, median, 75th percentile, and maximum degree (Table 2). Each individual graph was scored by how closely its own six statistics matched the per-generator averages. This yielded six per-statistic scores from 1 (best match) to 5000 (worst match), which were summed into a single ranking score for each graph. Graphs with lower overall ranking scores were thereby closer to the average degree distribution for their generator type. The 100 lowest-scoring graphs per graph type were retained on the basis that they should best represent the average degree distribution and would therefore produce disease simulations characteristic for that graph type.

**Table 2.**
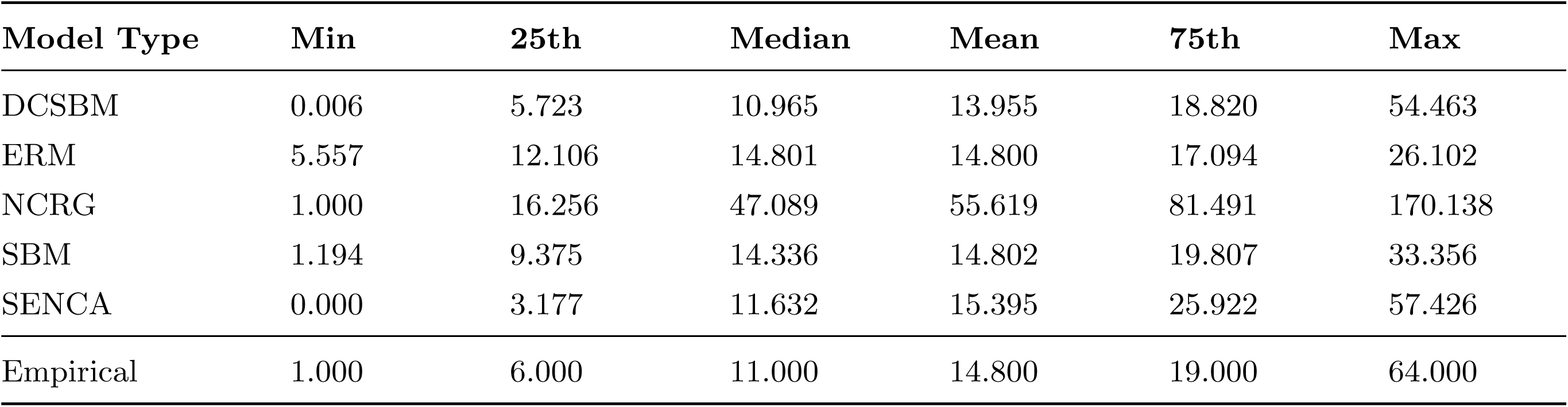
Degree summary statistics for each graph type averaged over 5000 graphs. For each graph per graph type, the minimum, 25th percentile, median, mean, 75th percentile, and maximum are calculated. These are then summarized by mean over all 5000 graphs (i.e., the minimum represents the average minimum degree for a specific graph type).

### Disease transmission models

Rabies transmission was simulated using an individual-based discrete stochastic susceptible-exposed-infectious-removed (SEIR) model, adapted from previously published models [3, 25]. The compartmental structure follows the standard susceptible-exposed-infectious-removed framework [2, 47]. The inclusion of exposed and removed compartments, combined with a declining susceptible population over time, is expected to slow disease progression and amplify the influence of heterogeneous contact structures. We additionally implemented a susceptible-infectious-susceptible (SIS) model, typically used to model for instance the common cold, as a comparative model in which individuals return directly to the susceptible pool upon recovery, thereby sustaining a larger susceptible population throughout the simulation. This allows assessment of how each graph type affects the overall susceptible population across varying infection periods and transmission rates representative of variation in SIS-infectious diseases. Model equations for the SEIR (Eq 1) and the SIS formulations (Eq 2) are given below; schematic representations, state variables and parameter descriptions are provided in the supplementary material (S4 Fig, S2 Table and S3 Table).

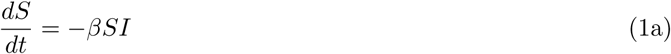

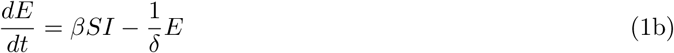

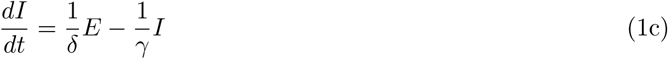

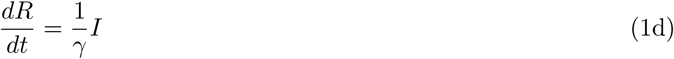

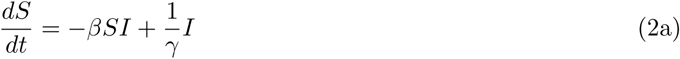

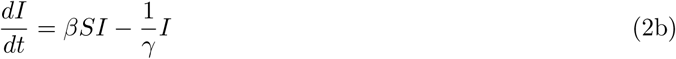

The SIS model simulation starts with a single infectious node *i*, randomly selected from the total set of vertices *V* of the network *G*(*V, E*) with *E* edges, such that *V_I_* (0) = *{i}* and *I*(0) = 1. The total number of infectious individuals at any time is represented by *I*(*t*) and *V_I_* (*t*) shows the set of vertices in the network that are infectious at time *t*. At each time step, the set of neighbors *N* (*V_I_* ) of *V_I_* (*t*) is identified, where the collective neighborhood of *V_I_* (*t*) is defined as the union of individual neighborhoods, *N* (*V_I_* ) = S_*i∈VI (t)*_ N(i) *N* (*i*) and is bound by the total population size (*|N* (*V_I_* )*| ≤ |V |*) in which *| · |* denotes the number of nodes in the enclosed set. For each *i*, the size of its individual neighborhood *N* (*i*) is dictated by its degree *k_i_*, where *k_i_ ≤ |V | −* 1. The state transition from susceptible to infectious is governed by a Bernoulli trial, which determines which susceptible neighbors *N_S_*(*i*) = *N* (*i*) *∩ V_S_*(*t*) will become infected, where *V_S_*(*t*) is the set of vertices that are susceptible at time *t*.

Specifically, each susceptible neighbor *j_S_ ∈ N_S_*(*i*) moves to the infectious state if *β ≥ r*, where *r ∈* [0, 1] is a pseudo-randomly generated number and *β ∈* [0.01, 0.2] is the transmission rate. The duration in days which the individual remains infected is sampled from a Poisson distribution with mean *λ* equal to the recovery period *γ*, such that *γ_i_* represents the infectious period of node *i*, and has a minimum duration of 1 day. We evaluate the model’s sensitivity across a range of *γ ∈* [3, 9] days. Individuals immediately return to the susceptible state when they are no longer infectious. The outbreak ends a) when there are no longer any individuals in the infectious class (*I*(*t*) = 0) or b) when the outbreak has reached its time limit at *t* = 1000.

The SEIR model follows the same process: the simulation starts with *V_I_* (0) = *{i}* and *I*(0) = 1, and a Bernoulli trial in which *P_infection_* = *β ≥ r* determines which of *i*’s susceptible neighbors *N_S_*(*i*) will be infected and therefore move to the exposed state, after which they become infectious. *E*(*t*) and *I*(*t*) detail the total number of exposed and infectious individuals at time *t* respectively. The duration in days during which individuals remain exposed and subsequently infectious are sampled from Poisson distributions with mean (*δ* = 30) for exposed individuals and mean (*γ* = 2) for infectious individuals, such that *δ_i_* and *γ_i_*represent the duration of the latent and infectious periods respectively for individual *i*. Once no longer infectious, individuals enter the removed state and exit the simulation, such that *R*(*t*) contains the total number of removed individuals at time *t*. The outbreak ends a) when there are no longer any individuals in the exposed and infectious class (*I*(*t*) + *E*(*t*) = 0), which may be a result of a failure to infect other nodes, a lack of susceptible neighbors (*N_S_*(*i*) = *∅*), or a combination of those two conditions, or b) when the outbreak has reached its time limit at *t* = 1000.

In the SEIR model, the average incubation period of 30 days was based on estimates reported in the literature on canine rabies. However, these vary widely across studies: while some report incubation periods as low as 14 days [48] and an average of approximately 18.6 days across mesocarnivorous reservoir hosts in a recent meta-analysis [49], others report higher values such as an average of 27.30 days during the 1948–1954 canine rabies epidemic in Tokyo [50], 21–60 days depending on rabies strain [51], and periods of up to 125 days [52]. We therefore opted for an average of 30 days modeled as a Poisson distribution, which has a standard deviation of *∼* 5.5 days and therefore captures most of the reported rabies incubation variation. While it excludes the extreme tail of incubation periods that has been recorded in a small minority of cases [52], this omission is unlikely to alter the comparative findings across generators reported here. The infectious period has a mean of two days because we assume that a symptomatic individual will be easily recognized as infected with rabies and hence killed as a result of cultural norms and practices before dying naturally from the disease. This assumption has been supported and successfully adopted in previous research [3, 53]. The SEIR infectious period was therefore held at this empirically established value, and *γ*-sensitivity was probed using the SIS model rather than by varying *γ* in the SEIR model, since longer SEIR infectious periods would no longer correspond to the majority of empirical scenarios.

We examined outbreak prevalence by calculating the average and peak number of daily active cases (*E*(*t*) + *I*(*t*)) in each outbreak. In these outbreaks, the total number of infections or cumulative incidence (*CI* ) was calculated as the sum of all unique infections throughout the entire outbreak duration. We additionally calculated the final outbreak size as the total fraction of removed individuals at the last day of each outbreak in the SEIR model. Outbreak duration was defined as the point at which the outbreak ceased due to exhaustion of the infectious and exposed classes or where the maximum time limit was reached.

We categorized outbreaks as minor or major based on the distribution of final outbreak sizes from SEIR simulations, using a kernel density estimate (*f* (*x*)) with a narrow bandwidth (*h* = 0.3) to preserve sensitivity to local features. The choice of *h* = 0.3 balances sensitivity to bimodality against smoothing of within-mode noise. Sensitivity analyses indicated that the distinction between minor and major outbreaks was robust to bandwidth adjustment: 98.7% of simulations retained the same classification across the full range of *h ∈* [0.1, 0.5]. Differences were concentrated among simulations with very low transmission (*β* = 0.01), with *≥* 99.97% of simulations remaining stable at all other Graph-*β* combinations. The classifications obtained with *h* = 0.2–0.3 were identical, supporting the use of *h* = 0.3 for the main analysis. A point was considered a peak if its height exceeded that of its two neighboring points and was at least 2% of the maximum density height. We calculated a dynamic threshold as the minimum value between the first and last significant local maxima, such that *τ*^ = arg min*_x∈_*_[*p*1_ *_,pL_*_]_ *f* (*x*), where *p*_1_ and *p_L_* denote the first and last peaks exceeding 2% of the maximum density height. This threshold was then used to categorize outbreaks as major or minor, and we summarized the mean and standard deviation of final outbreak size and outbreak duration for both categories.

### Simulations

We tested two parameter sampling approaches for the simulations: factorial and random sampling. Factorial sampling was initially used to thoroughly explore the entire parameter space, and its output was compared against that of random sampling to assess whether the latter could sufficiently cover the parameter space. Both approaches were used to run the disease transmission models for the empirical network and the synthetic graphs. For the factorial sampling approach, the design yielded a parameter grid of 5 (graph types) *×* 100 (graphs per graph type) *×* 39 (*β*) *× {*1 or 7*}* (*γ*) *×* 1000 (seeds), totaling 19,500,000 and 136,500,000 simulations for the SEIR and SIS models respectively (Table 3). The empirical graph was sampled with 3,900,000 and 27,300,000 parameter combinations respectively with unique seeds (1 *×* 39 (*β*) *× {*1 or 7*} γ ×* 1000 *×* 100), the latter for the 100 graphs to ensure equal sampling of the *β* and *γ* parameters (Table 3). Note that *δ*, fixed at a single value in the SEIR model, does not contribute additional combinations.

**Table 3.** Parameter combinations and sampling design for SEIR and SIS models. Numerical values for *β*, *γ*, and *δ* represent the number of discrete levels (steps) used in the grid rather than the specific parameter values or ranges (e.g., *δ* = 1 indicates a single value fixed at 30 days). For the factorial design, the total simulation count is the product of all parameter levels and repetitions. For the random sampling design, parameters are sampled independently for each outbreak. In the synthetic network scenarios, the total simulation volume is distributed evenly across the five graph types. Notably, there is a substantial difference in scale between the factorial and random sampling totals: factorial sampling produces orders of magnitude more simulations because every parameter combination is exhaustively crossed, while random sampling concentrates compute on a unique-seed-per-outbreak design.

| Approach | Model | Type | Index | $\beta$ | $\gamma$ | $\delta$ | Reps | Seeds | Total Simulations |
| --- | --- | --- | --- | --- | --- | --- | --- | --- | --- |
| Factorial | SEIR | Syn. | 500 | 39 | 1 | 1 | 1,000 | Unique | 19,500,000 |
|  |  | Emp. | 100 | 39 | 1 | 1 | 1,000 | Unique | 3,900,000 |
|  | SIS | Syn. | 500 | 39 | 7 | – | 1,000 | Unique | 136,500,000 |
|  |  | Emp. | 100 | 39 | 7 | – | 1,000 | Unique | 27,300,000 |
| Random | SEIR | Syn. | Rnd. | Rnd. | – | – | 1 | Unique | 2,500,000 |
|  |  | Emp. | 1 | Rnd. | – | – | 1 | Unique | 500,000 |
|  | SIS | Syn. | Rnd. | Rnd. | Rnd. | – | 1 | Unique | 5,000,000 |
|  |  | Emp. | 1 | Rnd. | Rnd. | – | 1 | Unique | 1,000,000 |

‘Rnd.’ indicates parameters sampled randomly from their respective distributions for each outbreak. ‘Unique’ seeds indicate that each simulation replicate was assigned a distinct random seed ( $s$ ).

While this approach is rigorous, it is time- and memory intensive, particularly for larger graphs with higher edge density. As a more efficient alternative, we set the total number of outbreaks per graph type at 500,000 for the SEIR model and 1,000,000 for the SIS model, given its larger number of potential parameter combinations, and randomly sampled a number of *γ* and *β* parameter configurations equal to the total outbreak number (Table 3). Each row of this parameter grid was then assigned a unique random seed. For synthetic graphs, rows were also assigned a random graph index between 1 and 100. A single ‘master’ seed (*s* = 100) was set prior to sampling to ensure reproducibility. Only outbreaks with at least one secondary infected individual were used for all summary statistics and visualizations, with the exception of the calculation of a threshold to distinguish between major and minor outbreaks in which outbreak simulations without a secondary infection were included.

Comparison of both approaches confirmed that random sampling sufficiently explored the parameter space and reproduced the patterns observed under the factorial approach. This approach was therefore used for all subsequent analyses reported in this manuscript and factorial sampling output is provided in S5 Fig–S7 Fig. All simulations were performed in R and were parallelized using the package *furrr* (v0.3.1).

## Results

### Network topology characterization

The five graph types produced distinct ensembles of stochastic networks (Fig 1), with variation in their distributions of network properties shown in Fig 2 (and S4 Table). Overall, graph types differ the least in their distributions of network density and mean degree, potentially because these two measures are not directly linked to network heterogeneity: edge density is only an indicator of mean graph connectedness. The log-normal shape of most empirical degree distributions results in mean degree being a poor indicator of heterogeneity.

**Fig 1.**
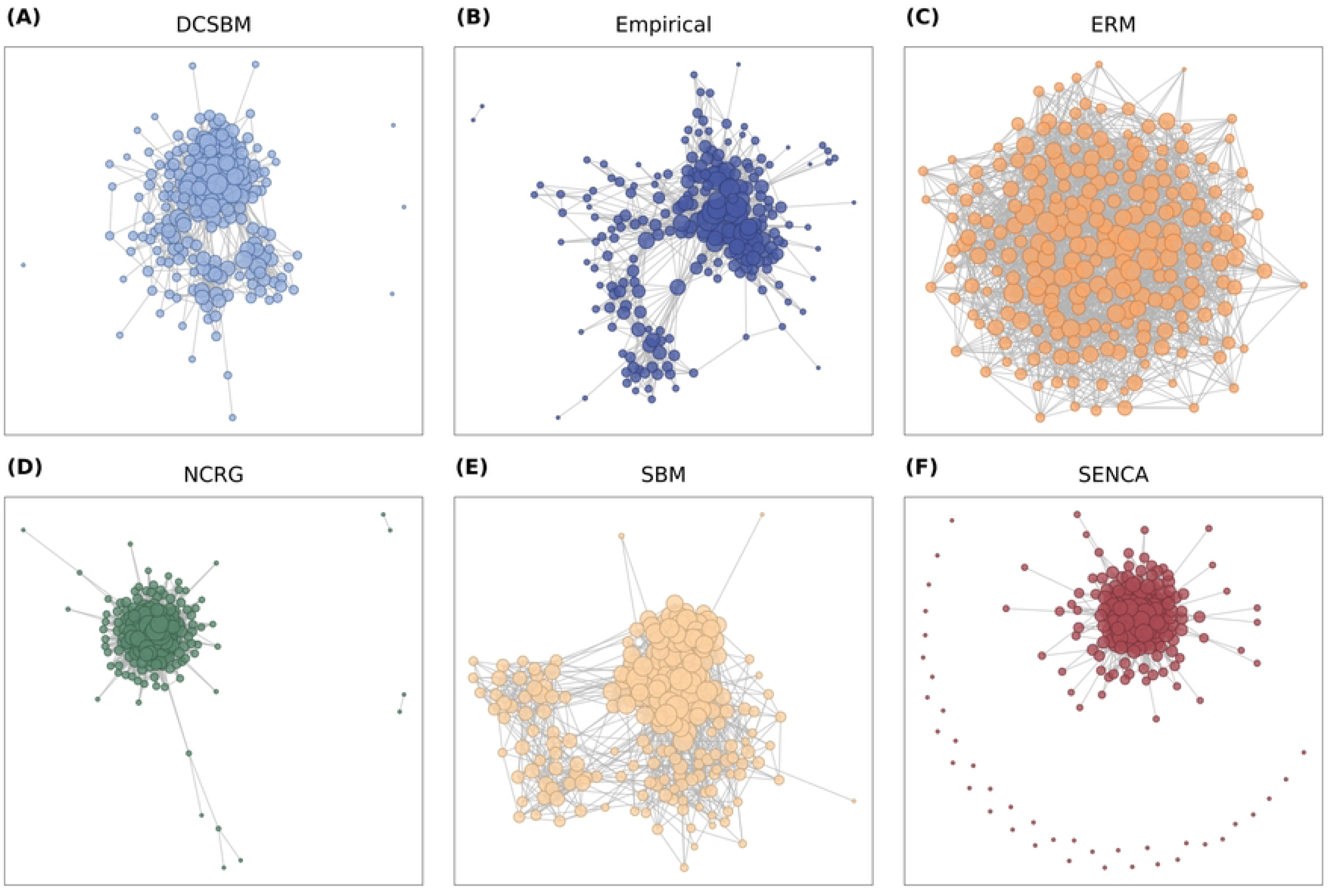
Example graph visualizations. A) DCSBM, B) Empirical network, C) ERM, D) NCRG, E) SBM, and F) SENCA. For each synthetic network, the graph whose degree distribution best fits the average degree summary statistics for that graph type is visualized in identical layout (see Methods). Nodes are sized as the square root of the node’s degree rescaled to a range between 0 and 10.

**Fig 2.**
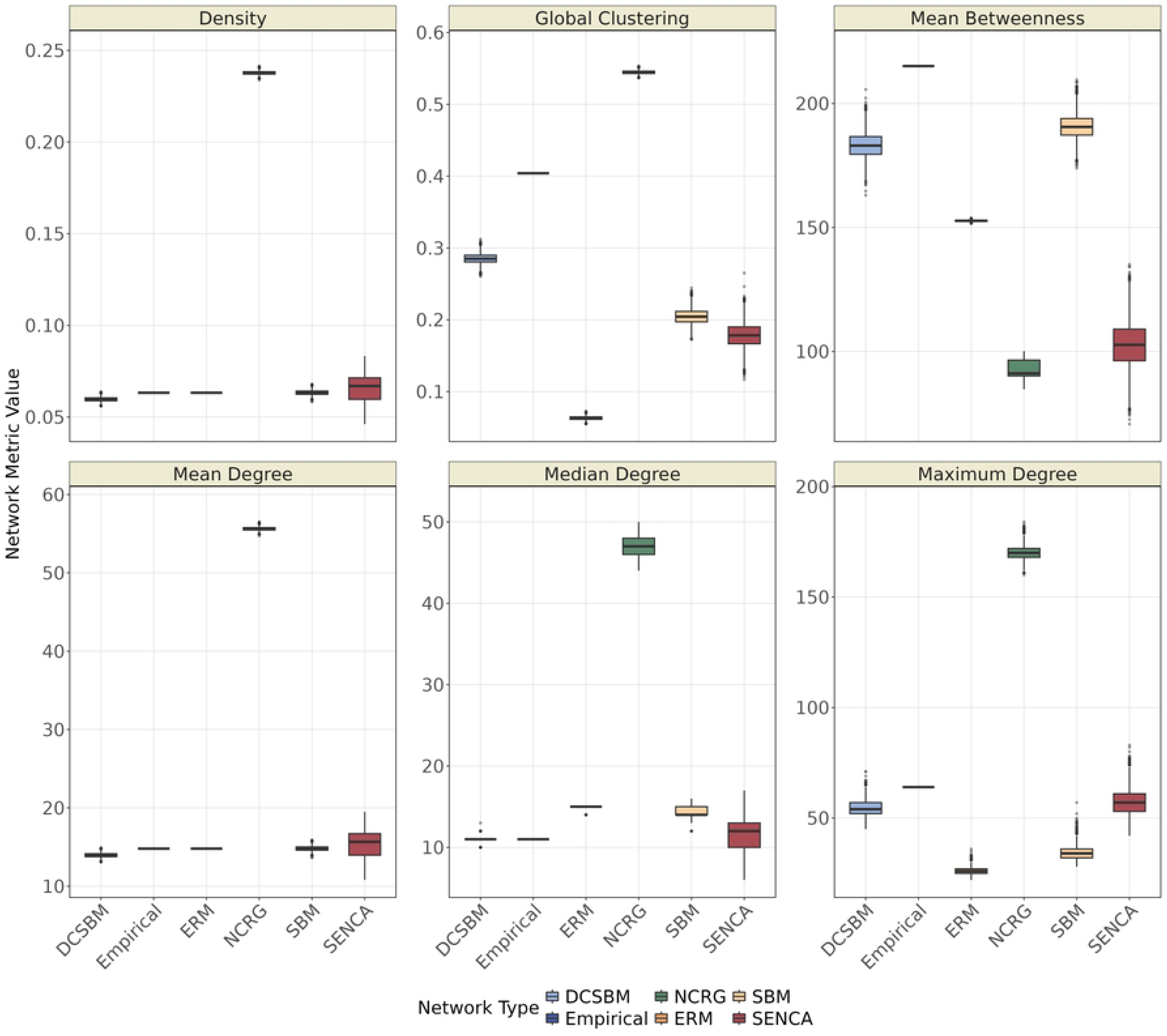
Network parameter distributions. Distributions of six network parameters — network density, global clustering, mean betweenness, mean degree, median degree, and maximum degree — over 5000 graph iterations of each graph type as well as their observed values in the empirical network. The six measures fall into two informative groups: density, mean degree, and median degree discriminate weakly among the generators because they are largely constrained by construction; global clustering, mean betweenness, and maximum degree discriminate strongly because they capture heterogeneity that only generators with explicit structural assumptions can reproduce.

The NCRG — which explicitly models triadic closure — has both a substantially higher mean degree (*µ* = 55.62) and edge density (*µ* = 0.24) compared to all other graph types and the empirical graph. This is likely because edges in the empirical network are often part of multiple triangles, which leads to a higher edge count and therefore also degree in the network reconstruction using the NCRG. The non-clustered random graph (ERM) generates a mean degree and network density identical to the empirical graph (*µ_degree_* = 14.80 and *µ_density_*= 0.06, respectively) as a result of its construction method. Of the remaining graph types, the SBM yields a mean degree (*µ_degree_* = 14.80) and network density (*µ_density_*= 0.06) most similar to the empirical graph in comparison to the other graph types. The SENCA produces most variation in these two measures (*µ_degree_* = 15.39 and *µ_density_* = 0.07 respectively), which is a pattern that is visible in its distributions of maximum and median degree as well (Fig 2). This is likely because it relies more heavily on stochastic sampling compared to the other generators, leading to more variation in its output graphs. The DCSBM produces similar mean edge density (*µ_density_*= 0.06) but lower mean degree (*µ_degree_* = 13.95) to the empirical graph and SBM. However, it fits closer to the empirical degree distribution’s tail: by better capturing the higher prevalence of low-degree nodes (below the 25th percentile), it more faithfully reproduces heterogeneity at the cost of a lower overall mean (Fig 3 and Table 2).

**Fig 3.**
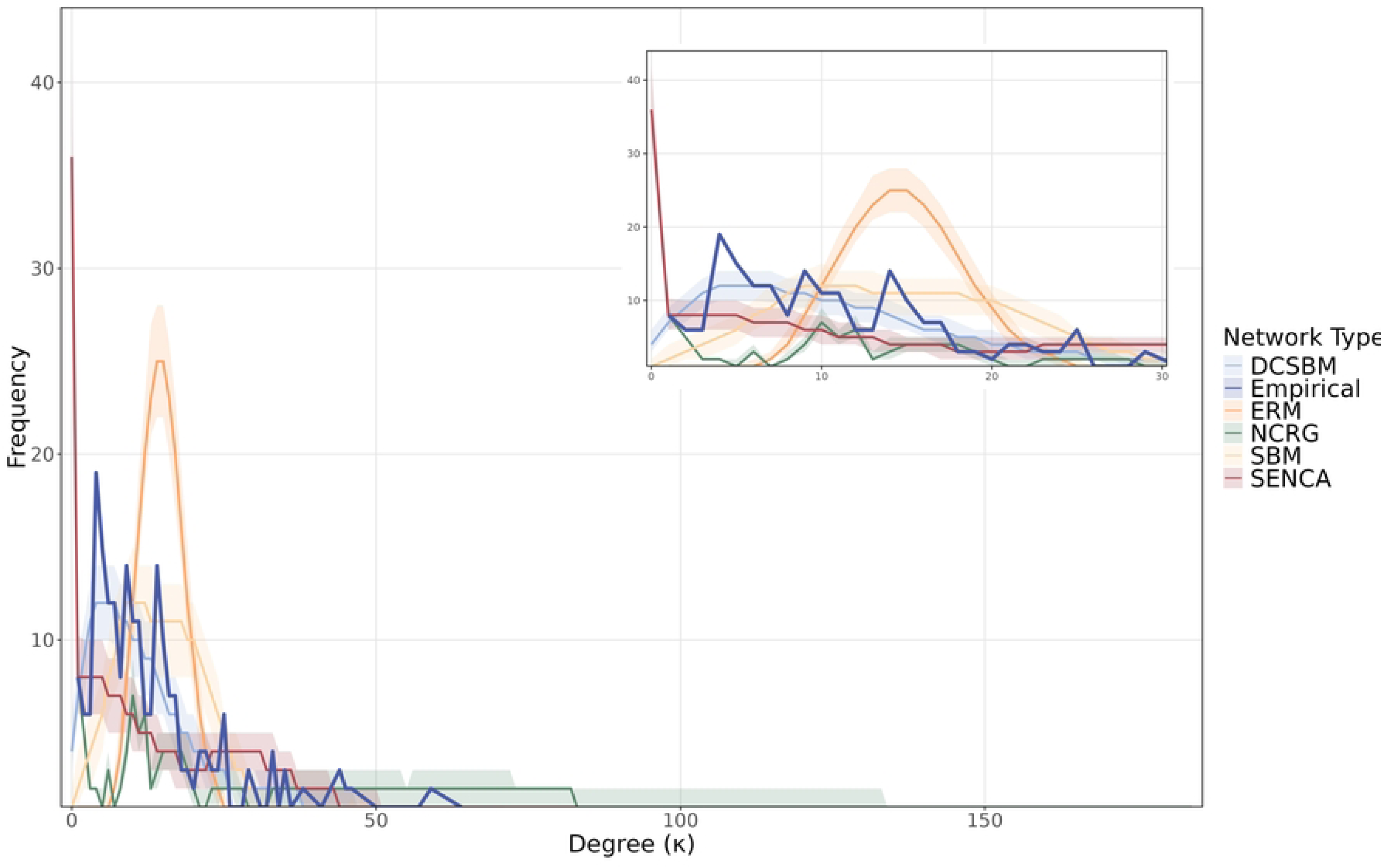
Summary of degree distributions across 5000 iterations of each graph type. Each point on the x-axis summarizes the distribution of frequency values observed across all graphs of a given graph type for a specific degree value. The shaded band indicates the interquartile range (25th to 75th percentile) and the solid line shows the median. The inset in the top right corner provides a zoomed-in perspective on the graph where the x-axis lies between 0 and 30.

In measures more indicative of heterogeneity, such as global clustering and mean betweenness, the graph types show more distinct variation in their distributions and deviate more clearly from the empirical network. With the exception of the NCRG, none of the graph types are able to match the empirical network’s relatively high clustering coefficient of 0.40. The empirical value as well as the NCRG’s higher clustering coefficient (*µ_clust_* = 0.54) support the idea that an individual’s neighbors are likely to be connected, leading to high clustering via triadic closure in empirical networks and the Newman clustered random graphs [54]. The DCSBM yields the second highest clustering coefficients (*µ_clust_* = 0.29), which lies closest to the empirical value when compared to all other graph types, followed by the SBM (*µ_clust_*= 0.20) and the SENCA (*µ_clust_* = 0.18). The higher value in the DCSBM compared to the SBM is likely a result of the heterogeneity in not only community membership, like the SBM, but also node connectivity, potentially leading to further clustering within and beyond communities. As expected, the mean clustering coefficient is lowest in the ERM (*µ_clust_* = 0.06), where triadic closure is comparatively rare [54].

No generator successfully reproduces the empirical mean betweenness of 215.1: it is highest in the two stochastic block models (*µ_betw_* = 190.7 and *µ_betw_* = 183.1 in the SBM and DCSBM respectively), followed by the ERM (*µ_betw_* = 152.7), the SENCA (*µ_betw_* = 102.6) and lastly the NCRG (*µ_betw_* = 92.1). The modeled community-structure in the DCSBM and SBM likely leads to the presence of scarce cross-community links which act as bridges between clusters and therefore strongly increase mean network betweenness (S8 Fig). In contrast, connections in the ERM are entirely independent and the shortest path between any two nodes can route through a large variety of alternative paths, keeping the mean and maximum betweenness centrality low (S8 Fig). Lower mean betweenness in the SENCA may be explained by the presence of many isolated individuals, whose low betweenness values negatively impact the network’s mean betweenness (Fig 1F). In the NCRG, the high density of edge triangles likely translates to a large number of alternative shortest paths and therefore lower betweenness scores for individual nodes.

The last two measures, median and maximum degree, give insight into the degree distribution of the synthetic graphs, because together they partially capture the expected log-normal shape of the empirical network’s degree distribution [55]. Here, the DCSBM and SENCA outperform the SBM, ERM and NCRG (Fig 2). This is to be expected, as these two generators explicitly model the degree distribution of the empirical network. The NCRG particularly overestimates median degree (*µ_med_* = 47.09) and maximum degree (*µ_max_* = 170.14), again likely linked to its overestimation of edges. The SBM and ERM both slightly overestimate median degree (*µ_med_*= 14.34 and *µ_med_*= 14.80 respectively) while underestimating maximum degree (*µ_max_* = 34.36 and *µ_max_* = 26.10 respectively). The SENCA generates an average median and maximum degree of 11.63 and 57.4 respectively, closely resembling the empirical observations of 11.0 and 64 respectively. However, this model shows considerable variation in its median degree and has a high number of maximum degree outliers that overestimate the maximum degree in relation to the empirical maximum degree by up to 11.6% compared to only 0.72% for the DCSBM. The DCSBM is overall much stabler in its performance and yields statistics (*µ_med_* = 10.97 and *µ_max_* = 54.5) that equally closely resemble the empirical network, but with more consistency and less variation across graphs in comparison to SENCA-generated graphs.

In addition, the DCSBM follows the shape of the empirical degree distribution most closely (Fig 3 and Table 2). As expected, the ERM and SBM follow a Poisson rather than log-normal degree distribution pattern, representing the homogeneity of this measure over the whole network (ERM) and within communities (SBM). The SENCA produces a higher number of individuals with zero connections, likely because edges are initially created based on the scaled Euclidean distance between two nodes. As a result, there might be a higher-than-expected number of individuals who are far removed from others and hence have no neighbors. In contrast, the NCRG underestimates the number of individuals with few connections, but yields a thicker and longer tail with degrees not observed in the empirical network. Kolmogorov-Smirnov distances between the synthetic and empirical degree distributions further confirm this pattern (S3 Fig).

Overall, these data imply that while the DCSBM generator does not perform best in each separate network measure, it does appear to best capture both global (clustering) and local (degree and betweenness) heterogeneity in the empirical network.

Comparing the performance of the graphs across all measures, we found that the DCSBM yielded the lowest Mahalanobis distance of 17.57, followed by the SBM with 20.98 and the SENCA with 29.21. The NCRG and ERM differ most, yielding distances of respectively 188.74 and 220.67. These gaps between generator means are large relative to within-generator variance, as visible in the per-measure distributions of Fig 2, so the ranking is unlikely to be a sampling artifact. While the ERM’s extreme statistic may be due to its lack of clustering and its strongly deviating degree distribution, the NCRG appears to over-correct for clustering and therefore inflates the empirical degree distribution. Taken together, the DCSBM graph generator performs best in reconstructing the empirical network’s overall structural network properties and capturing its heterogeneity.

### Epidemic dynamics across network topologies

In the SEIR simulation (*γ* = 2), the time-averaged prevalence (*Ī*) and peak prevalence (*I_max_*) largely exhibit consistent trajectories across most graph types. In the empirical network outbreaks, low transmission rates (0 *< β ≤* 0.03) generate near-zero average and peak prevalence. This is followed by a sharp nonlinear inflection between *β* = 0.03 and *β* = 0.07, marking the onset of major outbreaks. At higher transmission rates, growth decelerates as the shrinking pool of susceptible individuals leads to network saturation (Fig 4A).

**Fig 4.**
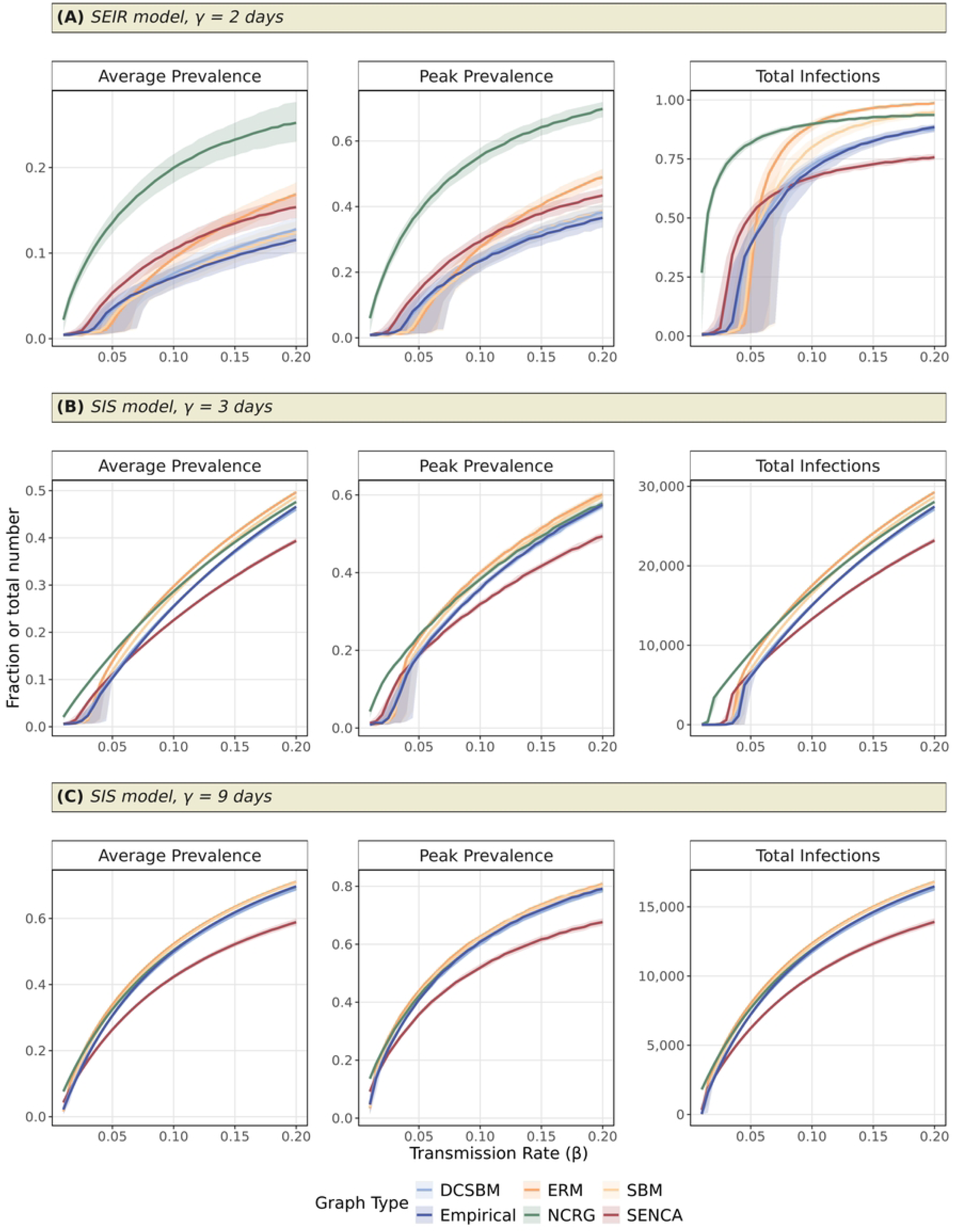
Outbreak simulation outcomes for SEIR and SIS models. Outbreak dynamics are illustrated for the A) SEIR model with *γ* = 2, B) SIS model with *γ* = 3, and C) SIS model with *γ* = 9 using the average prevalence (*Ī*), the peak prevalence (*I_max_*) and the number of total infections (*CI*, as a fraction in A but count in B and C) for each of the five graph types in comparison to the empirical network across the full range of modeled transmission rates. Only outbreaks with at least one secondary infection are shown in this visualization.

The SBM and DCSBM configurations most closely capture the empirical network’s outbreak dynamics. The SBM exhibits a minor delay in reaching the epidemic threshold until *β* = 0.07, but then rapidly converges with the DCSBM and empirical network (Fig 4A). The DCSBM captures the empirical average and peak prevalence very well, despite slight inflation for *β >* 0.08 for average prevalence and *β >* 0.13–0.14 for peak prevalence. The ERM displays a minor delay, but subsequently increases more sharply beyond *β* = 0.07. This accelerated transmission is likely driven by the model’s generation of short path lengths, which facilitate rapid transmission throughout the network in the absence of local clustering constraints. Conversely, the SENCA consistently overestimates outbreak prevalence across all transmission rates. The higher number of active cases in this model is an expected consequence of its higher average degree and lack of local clustering, both of which remove barriers to transmission provided the initially infected node is not isolated. The NCRG deviates more from these patterns. At a near-zero transmission rate (*β* = 0.01), its high edge density and triangular structures circumvent the typical threshold bottleneck. It consistently maintains an approximately 10–13% higher average and peak prevalence in comparison to the other graph types, which can likely be attributed to its comparatively high number of transmission paths.

The empirical distribution of total number of unique infections (*CI* ) is characterized by a transmission threshold at around *β* = 0.03 with a median CI (*Med_CI_* ) of 0.021, followed by exponential growth up to *β* = 0.07 (*Med_CI_* = 0.545) after which it gradually slows to the point where higher transmission rates exhaust the majority of the available susceptible population (for *β* = 0.20, *Med_CI_* = 0.885) (Fig 4A). While the other graph types generate a similar distribution shape, the NCRG again shows exponential growth even for low transmission rates, yielding median outbreak size of 72.8% for *β* = 0.03. This surge stabilizes around *β* = 0.07 and does not increase much further for *β* = 0.20 (*Med_CI_* = 0.936). The SENCA also initially rises faster, with a median 18.7% of the population having been infected for *β* = 0.03, but slows down for higher transmission rates (*Med_CI_* = 0.757 for *β* = 0.20). In contrast, the ERM has an initially delayed response (*Med_CI_* = 0.013 for *β* = 0.03), followed by a drastic surge which continues to increase steadily and reaches nearly the entire population at *β* = 0.20 (*Med_CI_* = 0.987). The SBM follows a fairly similar trajectory with an initial underestimation (*Med_CI_* = 0.013 for *β* = 0.03), but inflates the empirical outbreak pattern for transmission rates higher than 0.05, ultimately reaching 94.9% of the population for *β* = 0.20. The DCSBM best fits the empirical distribution for both lower (at *β* = 0.03, *Med_CI_* = 0.021) and higher transmission rates (at *β* = 0.20, *Med_CI_* = 0.876), despite its slight overestimation in outbreak size of around 2% for 0.06 *< β ≤* 0.13.

Comparing the SIS simulation with its lowest and highest *γ*values to the SEIR simulation outbreaks, it is clear that the SIS model yields smaller differences in outbreak size across transmission rates regardless of the length of the infectious window (Fig 4B–C and S9 Fig for remaining *γ* values). For *γ* = 3, the ERM, SBM, and DCSBM all closely follow the empirical outbreak trajectory in all three output measures, with the DCSBM nearly perfectly following the empirical network. The NCRG shows a general overestimation of outbreak size, which is initially more pronounced but gradually decreases to around 1% or less for *β* = 0.20. The SENCA also initially slightly overestimates prevalence and cumulative incidence for *β <* 0.05, but underestimates them for all *β >* 0.05 likely due to presence of isolated individuals. While the SBM and ERM follow the empirical outbreak trajectory, they inflate cases and infections by 1–2%, which again may be attributable to the higher average and median degree of individuals in these networks.

At *γ* = 9, the longer infectious period leads to a higher number of individuals being infected at any given time while simultaneously resulting in a lower total number of infections as the turnover of individuals is slower (Fig 4C). For this longer infectious period, the convergence between graph types is even higher. The SBM, ERM, NCRG, and DCSBM follow the empirical case pattern very well (for *β* = 0.20, *Med_Ī_* is 0.708, 0.711, 0.691 and 0.688 respectively compared to 0.697 in the empirical network). The SENCA is an exception to this trend; it again consistently and increasingly underestimates cases for higher transmission rates (for *β* = 0.20, *Med_Ī_* = 0.589). Hence, while the DCSBM remains among the closest to the empirical network across the full transmission rate range, this convergence likely reflects how heterogeneous network structure is no longer as crucial for transmission in a model without a removal state and with longer infectious periods. This may therefore lead to a homogenization of the case pattern and remove the influence of network hubs and super-spreader individuals.

### Outbreak classification and scale across network topologies

The observed outbreak size patterns are inextricably linked to the outbreak duration too, particularly in the SEIR model where all outbreaks ended prior to reaching the maximum outbreak duration (Fig 5). We distinguished between minor and major outbreaks for each graph type and *β* combination by identifying local peaks and their corresponding valleys (Materials and methods and S10 Fig). At *β* = 0.01, most graph types closely mirror the empirical network, with major outbreaks averaging 1.6–2.3% of the population over 69–87 days and minor outbreaks remaining near-negligible (*µ_size_ ≈* 0.5%*, µ_duration_ ≈* 4.5 days). The NCRG is a clear exception: its major outbreaks already average 28.1*±*15.2% of the population and last 337.2*±*154.9 days on average, reflecting the same threshold-bypassing pattern observed in the prevalence analysis driven by its high edge density and abundance of triangular structures (S5 Table).

**Fig 5.**
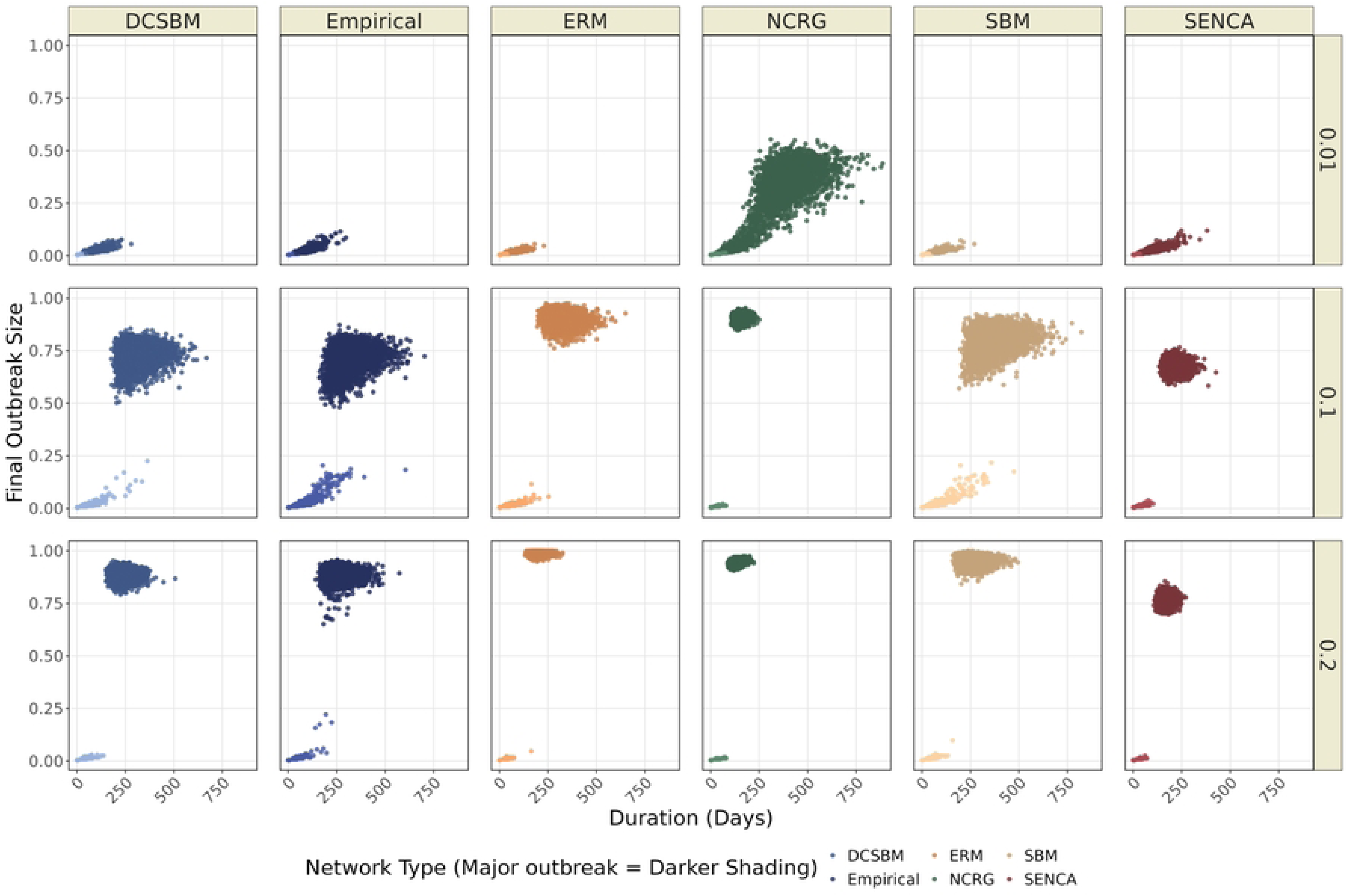
SEIR final outbreak size and outbreak duration. Results are shown for three values of *β* representing its full modeled range, separated into minor and major outbreaks for outbreaks simulated with each graph type and the empirical network. The x-axis shows outbreak duration measured in days (maximum = 1000) and the y-axis shows the final outbreak size measured as the proportion of removed individuals at the final day of the outbreak. Major outbreaks are indicated with darker shading compared to minor outbreaks.

At *β* = 0.1, the empirical network separates into two distinct clusters, with major outbreaks containing on average 71.5*±*5.3% of the population and lasting on average 319.9*±*68.6 days (S10 Fig and S5 Table). The DCSBM and SBM reproduce this pattern most closely (average major outbreak size: 73.1*±*4.4% and 80.5*±*5.1%; average duration: 311.9*±*61.2 and 362.9*±*77.0 days respectively), with the SBM exhibiting slightly more size variability and a higher maximum final outbreak size (92.3% versus 85.5%). The SENCA produces two clusters that are more concentrated, yielding a slightly lower average size of 67.8*±*2.5% and substantially shorter average duration of 213.0*±*31.3 days consistent with earlier susceptible population exhaustion. The NCRG exhibits more divergence between minor and major outbreaks. On average, its outbreaks are larger (90.3*±*1.4%) but of substantially shorter and less variable length (149.6*±*18.5 days).

Lastly, the ERM shows a similar gap between minor and major outbreaks (89.9*±*2.8%) and a more comparable duration (313.4*±*51.3 days) to the empirical scenario (S5 Table). At *β* = 0.2, clusters concentrate further: empirical outbreaks average 88.7*±*2.6% of the population over 254.9*±*45.0 days, while minor outbreaks average 0.5*±*0.5% of the population over 3.8*±*12.0 days (S5 Table). The DCSBM again fits most closely (major outbreaks: *µ_size_* = 88.3 *±* 2.1%, *µ_duration_* = 227.9 *±* 34.7 days; minor outbreaks: *µ_size_* = 0.5 *±* 0.2%, *µ_duration_*= 2.9 *±* 8.5 days), but yields a more concentrated distribution of values (S10 Fig and S5 Table). The SBM also captures the empirical patterns well, though with slightly higher major outbreak sizes (*µ_size_*= 95.2 *±* 1.9%, *µ_duration_*= 254.9 *±* 41.1 days) and shorter minor outbreaks (*µ_size_* = 0.5 *±* 0.2%, *µ_duration_* = 2.9 *±* 9.2 days). The SENCA overall produces shorter, smaller and less variable outbreaks of both types (major outbreaks: *µ_size_*= 76.2 *±* 2.0%, *µ_duration_*= 163.2 *±* 20.5 days; minor outbreaks: *µ_size_* = 0.4 *±* 0.1%, *µ_duration_*= 2.2 *±* 5.2 days). The ERM generates major outbreaks that encompass nearly the entire population and are most homogeneous across iterations (*µ_size_*= 98.9 *±* 0.7%, *µ_duration_*= 193.8 *±* 23.5 days) and generates relatively small and short minor outbreaks (*µ_size_* = 0.4 *±* 0.1%, *µ_duration_* = 1.9 *±* 6.0 days). Lastly, the NCRG portrays a similar pattern as for *β* = 0.1, but with larger major outbreaks (*µ_size_* = 94.2 *±* 0.9%, *µ_duration_* = 123.6 *±* 17.6 days) and comparably similar minor outbreaks (*µ_size_* = 0.4 *±* 0.1%, *µ_duration_* = 2.0 *±* 5.9 days).

Across all these transmission rates, the DCSBM most consistently reproduces the empirical distribution of outbreak sizes and durations. The SBM performs comparably to the DCSBM but inflates major outbreak size, particularly at *β* = 0.2. The ERM and NCRG both overestimate major outbreak saturation, particularly at a higher transmission rate, while the SENCA systematically underestimates outbreak duration, likely reflecting the structural constraints discussed above.

### Generalizability across empirical and simulated scenarios

To test whether the patterns found so far are network-specific, we compared four additional proximity-based dog interaction networks of varying sizes, number of connected components, and degrees of clustering (S11 Fig). The Romana and Hepang networks contain fewer nodes compared to the empirical (Chad), Sabaneta and Habi networks, while these three networks are characterized by lower edge density (Table 4). Moreover, the Habi and Romana networks are both fully connected, whereas the Hepang and Sabaneta networks each contain one major component encompassing nearly all nodes and a smaller isolated cluster, similar to the thus far examined Chad network.

**Table 4.** Empirical network metrics, synthetic-model medians and Mahalanobis distance (*D*^2^) across the five empirical locations. For each location, the table reports median values across 5000 graph iterations for six network measures (density, global clustering, mean betweenness, mean degree, median degree, maximum degree) for each generator alongside the empirical value, and the Mahalanobis distance summarizing the multivariate gap between each generator’s measure distribution and the empirical reference.

| Location | Parameter | Empirical | ERM | SBM | DCSBM | NCRG | SENCA |
| --- | --- | --- | --- | --- | --- | --- | --- |
| **Chad**<br>(Nodes: 235)<br>(Edges: 1739) | Density | 0.063 | 0.063 | 0.063 | 0.060 | 0.238 | 0.067 |
|  | Global Clustering | 0.404 | 0.063 | 0.204 | 0.285 | 0.545 | 0.178 |
|  | Mean Betweenness | 215.089 | 152.681 | 190.555 | 183.026 | 91.162 | 102.653 |
|  | Mean Degree | 14.800 | 14.800 | 14.804 | 13.949 | 55.617 | 15.664 |
|  | Median Degree | 11.000 | 15.000 | 14.000 | 11.000 | 47.000 | 12.000 |
|  | Max. Degree | 64.000 | 26.000 | 34.000 | 54.000 | 170.000 | 57.000 |
|  | *Mahalanobis $D^2$* | — | 220.673 | 20.984 | 17.570 | 188.741 | 29.208 |
| **Romana**<br>(Nodes: 57)<br>(Edges: 275) | Density | 0.172 | 0.172 | 0.172 | 0.164 | 0.393 | 0.085 |
|  | Global Clustering | 0.392 | 0.172 | 0.197 | 0.299 | 0.597 | 0.193 |
|  | Mean Betweenness | 29.825 | 27.526 | 28.246 | 29.912 | 15.456 | 31.333 |
|  | Mean Degree | 9.649 | 9.649 | 9.649 | 9.193 | 22.000 | 4.772 |
|  | Median Degree | 8.000 | 10.000 | 10.000 | 8.000 | 22.000 | 4.000 |
|  | Max. Degree | 27.000 | 16.000 | 16.000 | 24.000 | 44.000 | 15.000 |
|  | *Mahalanobis $D^2$* | — | 18.963 | 15.790 | 5.277 | 67.896 | 4.731 |
| **Sabaneta**<br>(Nodes: 121)<br>(Edges: 471) | Density | 0.065 | 0.065 | 0.065 | 0.063 | 0.165 | 0.098 |
|  | Global Clustering | 0.378 | 0.064 | 0.185 | 0.231 | 0.375 | 0.177 |
|  | Mean Betweenness | 121.182 | 92.802 | 117.273 | 116.223 | 65.934 | 76.963 |
|  | Mean Degree | 7.785 | 7.785 | 7.785 | 7.562 | 19.835 | 11.719 |
|  | Median Degree | 7.000 | 8.000 | 8.000 | 7.000 | 18.000 | 11.000 |
|  | Max. Degree | 27.000 | 15.000 | 17.000 | 26.000 | 72.000 | 28.000 |
|  | *Mahalanobis $D^2$* | — | 53.696 | 14.771 | 9.791 | 74.238 | 20.624 |
| **Habi**<br>(Nodes: 81)<br>(Edges: 176) | Density | 0.054 | 0.054 | 0.054 | 0.050 | 0.098 | 0.158 |
|  | Global Clustering | 0.419 | 0.053 | 0.191 | 0.271 | 0.346 | 0.235 |
|  | Mean Betweenness | 120.593 | 82.185 | 98.235 | 84.580 | 59.821 | 41.000 |
|  | Mean Degree | 4.346 | 4.346 | 4.321 | 4.025 | 7.827 | 12.667 |
|  | Median Degree | 4.000 | 4.000 | 4.000 | 3.000 | 5.000 | 12.000 |
|  | Max. Degree | 14.000 | 10.000 | 10.000 | 14.000 | 27.000 | 27.000 |
|  | *Mahalanobis $D^2$* | — | 27.967 | 7.514 | 5.741 | 30.210 | 19.311 |
| **Hepang**<br>(Nodes: 61)<br>(Edges: 217) | Density | 0.119 | 0.119 | 0.119 | 0.105 | 0.240 | 0.081 |
|  | Global Clustering | 0.463 | 0.117 | 0.297 | 0.380 | 0.533 | 0.193 |
|  | Mean Betweenness | 43.377 | 38.311 | 53.951 | 49.246 | 23.705 | 34.967 |
|  | Mean Degree | 7.115 | 7.115 | 7.115 | 6.295 | 14.393 | 4.885 |
|  | Median Degree | 6.000 | 7.000 | 7.000 | 5.000 | 11.000 | 4.000 |
|  | Max. Degree | 34.000 | 13.000 | 15.000 | 25.000 | 41.000 | 16.000 |
|  | *Mahalanobis $D^2$* | — | 27.536 | 14.865 | 5.519 | 36.020 | 9.917 |

Examining the network topology for each of the four networks, the DCSBM overall best captures empirical maximum and median degree and most closely reproduces each network’s overall degree distribution, yielding the lowest K-S distances to empirical degree distributions among all graph types (Table 4 and S12 Fig–S14 Fig). The ERM and SBM produce similar mean and median degree outcomes. The SENCA varies in its degree outcomes, underestimating degree in the smaller Hepang and Romana networks, while overestimating it in the larger but lower-density Sabaneta and Habi where it also underestimates betweenness and clustering. This effect is most severe for Habi, the lowest density network. The NCRG consistently overestimates degree statistics, particularly median and maximum degree in Sabaneta and edge density in Hepang and Romana. With the exception of average betweenness, it performs best in reproducing Habi’s topology in comparison to the other networks. These patterns reflect the same tendency seen in the Chad network to generate triangular structures and high-degree hubs regardless of the empirical degree distribution. In comparison to degree measures, mean betweenness and global clustering vary more across graph types.

The synthetic Romana networks yield narrow mean betweenness distributions closely resembling the empirical network, but for Habi and Sabaneta they mostly underestimate mean betweenness. The SBM yields the highest mean betweenness overall, while the NCRG, SENCA and ERM produce lower values, particularly in less dense networks. The SBM, SENCA and ERM strongly underestimate global clustering. The DCSBM also systematically underestimates global clustering, but yields estimates far closer to the empirical values, competing with the NCRG for better performance between network locations. The NCRG does not produce such a consistent pattern, overestimating clustering for some and underestimating it — albeit only slightly — for others. Its higher performance for this metric is no surprise, as the model is designed to capture this metric. Still, the DCSBM is able to yield nearly comparable clustering without optimizing for it. In combination with its consistent improvement over the SBM and the inability of the other synthetic graphs to reproduce clustering, this suggests that a) clustering does not need to be specifically targeted for realistic clustering estimates to emerge and b) degree heterogeneity plays an important role in modeling empirical transitivity patterns. This conclusion is further supported by the DCSBM’s lower Mahalanobis distance to the empirical network’s combined summary measures at every location except Romana (Table 4).

These structural differences impact outbreak dynamics. The two lower density networks (Habi and Sabaneta) produce simulated empirical rabies outbreaks with lower average and peak prevalence than Hepang and Romana (Fig 6). The ERM, SBM, and DCSBM largely follow the simulated outcomes of the empirical network, particularly for average and peak prevalence measures. The NCRG and SENCA diverge more, where the NCRG consistently overestimates all outbreak metrics when compared to the empirical network, only coming close to empirical estimates of total infections (*CI*) in the Romana network for larger transmission rates. The SENCA yields more variable outcomes, matching the NCRG’s overestimation for the Habi and Sabaneta networks while underestimating prevalence for the Romana network. Interestingly, it closely matches Hepang’s empirical outbreak dynamics. These variable outcomes are likely attributable to how the SENCA models edge density in these networks: it overestimates edge density in the former two networks, yielding more transmission pathways, while underestimating it for the Romana and Hepang networks.

**Fig 6.**
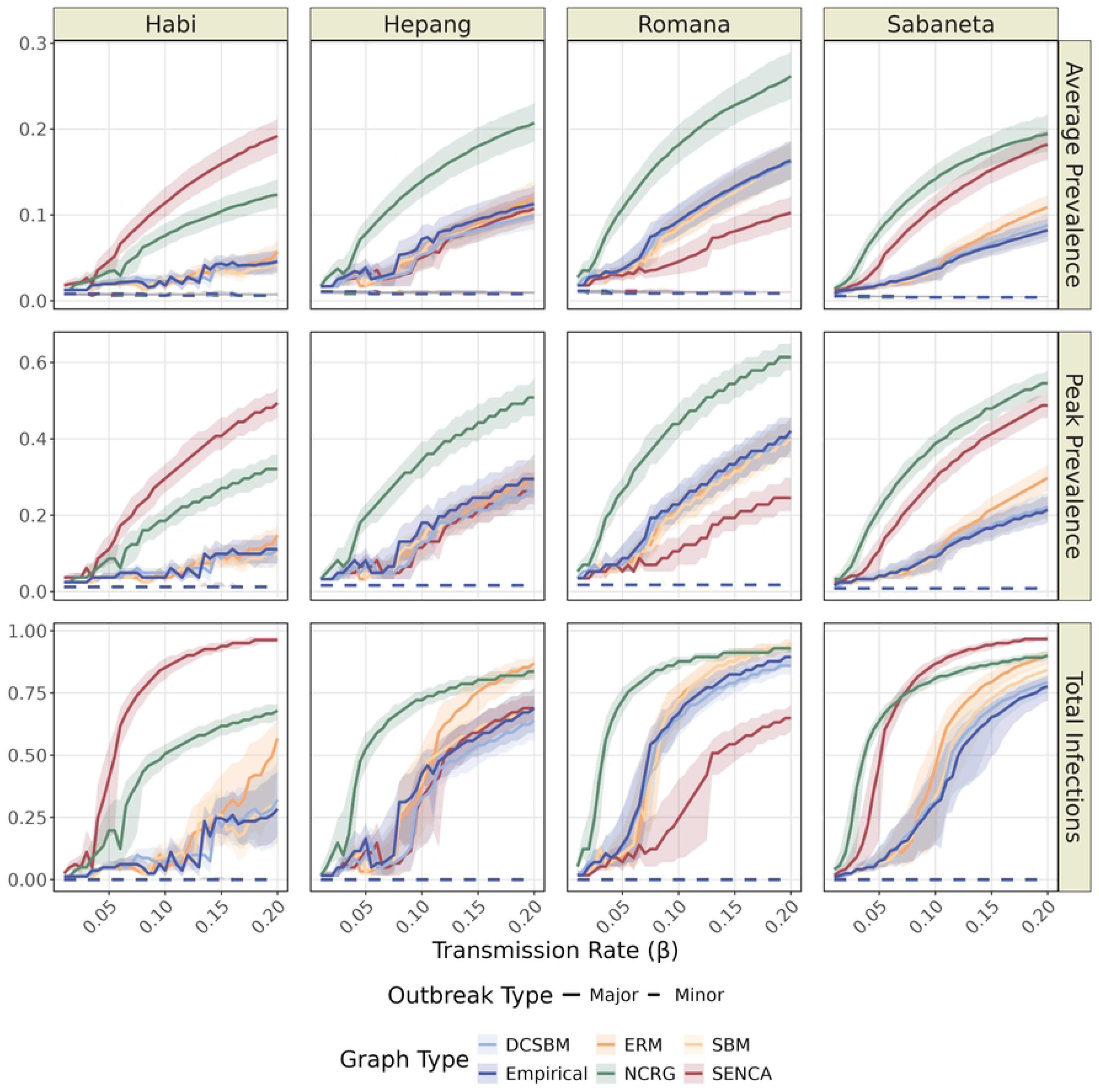
SEIR outbreak dynamics for each location per graph type. Panels visualize the relationship between transmission rate and average and peak prevalence as well as the total infections (*CI*) for each location network, separated into minor (dashed line) and major (solid line) outbreaks.

Overall, the ERM does an acceptable job at reproducing outbreak dynamics, though it inflates *CI* for high transmission rates in all four networks. It yields considerably higher *CI* in the Habi, Sabaneta and Hepang networks for *β >* 0.15, most likely driven by its short path lengths in the absence of modeled clustering. The SBM and DCSBM perform consistently well and produce comparable outbreak dynamics outcomes, portraying only negligible differences in favor of the DCSBM. Together, these findings highlight how the DCSBM consistently reproduces both empirical network topology and outbreak dynamics, thereby supporting its applicability beyond a single network.

## Discussion

In this study, we evaluated five network generator algorithms for network reconstruction in rabies transmission modeling, demonstrating how structural choices in network reconstruction can alter outbreak predictions as substantially as the biological parameters of the disease model itself. Across both static topological metrics and stochastic outbreak simulations, the degree-corrected stochastic block model was the most consistent at reconstructing free-roaming dog contact networks and reproducing their empirical rabies transmission dynamics. Crucially, this consistency was robust across five distinct empirical networks of varying size, density, and connectedness.

Contrasting the performance of the DCSBM against the other four network generators reveals the relative contributions of three fundamental axes of network heterogeneity: degree heterogeneity, community structure, and local clustering. Comparing the DCSBM to the standard SBM demonstrates how modeling within-block degree heterogeneity on top of community structure improves both topological and epidemiological fit.

Conversely, the contrast between the DCSBM and SENCA illustrates the substantial explanatory value of community structure beyond degree heterogeneity alone. The comparison with the NCRG introduces a third axis: explicit triadic closure, which the DCSBM does not model natively. Although local clustering is a well-documented empirical feature of interaction networks, network reconstruction based on triadic membership alone did not outperform the implicit clustering that naturally emerges from combining degree heterogeneity with community structure within the DCSBM framework. The NCRG produced the most pronounced deviations from empirical outbreak dynamics, a direct consequence of its high edge density and abundance of triangular structures generating transmission pathways that circumvent the barriers present in the empirical network.

The NCRG outcome suggests a more general working hypothesis worth testing in future work: that the three axes of degree heterogeneity, community structure, and clustering do not decompose orthogonally as independent reconstruction targets. The NCRG isolates clustering and produces a roughly fourfold edge inflation relative to the empirical network (mean degree 55.6 vs empirical 14.8; density 0.24 vs empirical 0.06), and the implicit clustering that emerges from the DCSBM’s joint coupling of degree heterogeneity with community structure (mean global clustering 0.29) is closer to the empirical value (0.40) than the NCRG’s explicit target-clustering value (0.54). This pattern is consistent across all five empirical locations examined here (Table 4), where the DCSBM achieves Mahalanobis distances several-fold smaller than the NCRG’s at every location, though the NCRG yields slightly more accurate clustering estimates for 3 out of 5 locations. We frame this as a working hypothesis rather than an established principle because our evidence rests on a single network family (free-roaming dog proximity contacts in five geographic settings), and the joint-stub formulation we implemented is one of several possible NCRG-class constructions. If the hypothesis is borne out by independent testing in other empirical network families, it would imply that orthogonal-axes generator-comparison frameworks materially understate the difficulty of network reconstruction, and that practical generator selection should weight joint structural coupling more heavily than independent feature targeting.

Importantly, the influence of these structural choices is not static; rather, it is highly dependent on the life-history parameters of the disease model. While degree heterogeneity and community structure are recognized as foundational drivers of outbreak dynamics and intervention design [11, 16, 21], our SIS sensitivity analyses with varying infectious periods reveal that this topological influence progressively weakens as the infectious window lengthens. At a nine-day infectious period, average and peak prevalence trajectories converged across nearly all generators (*Ī ≈* 0.69 to 0.71 at *β* = 0.20, all within roughly two percentage points of the empirical value of 0.697), even as their underlying topological signatures remained clearly distinct. The SENCA remained the sole exception to this convergence, likely reflecting the influence of its smaller fully connected component.

This convergence highlights how the extent to which contact network topology shapes outbreak dynamics depends on how the pathogen exploits that network. When infectious windows are short relative to typical contact rates — as in canine rabies, where the symptomatic period averages only a few days due to human recognition of behavioral symptoms and subsequent euthanasia — each infected individual can only make a limited number of transmission attempts. Under this assumption, the exact topological identity of those immediate contacts matters intensely. As the infectious window lengthens, however, infected individuals progressively exhaust their local network neighborhoods, and the precise ordering of contacts becomes less determinative of overall outbreak trajectory. Network reconstruction therefore appears most consequential for short-infectious-window pathogens like canine rabies, and progressively less so for slower-burning infections in which the pathogen has sufficient time to traverse the giant connected component regardless of local structure and bottlenecks. This pattern aligns with prior theoretical work on the relationship between infectious-window length and topological sensitivity [2, 19], and our results suggest the effect is large enough to inform generator selection in practical modeling applications. These implications extend past canine rabies to other short-infectious-window infectious diseases where network-based control strategies are actively considered.

Several limitations bound these findings. The first considers spatial scale. All five empirical networks contain between 57 and 235 nodes; whether the generator rankings established at this scale hold for city-scale populations of thousands of dogs — the regime relevant to large-scale mass vaccination programs — remains untested. Stochastic block model inference becomes computationally demanding at that scale, and triangle-based generators face parallel scaling constraints. A second limitation involves network sampling. While the spatiotemporal tracking of contact networks has advanced via GPS and other real-time location systems through increased positional accuracy and shorter temporal intervals [56–59], empirical data remains restricted to a participating or captured subset of the (animal) population. This introduces an inevitable trade-off between sampling coverage, duration, and frequency [1, 3, 28, 60–62], leading to restrictions in estimates of epidemic outbreaks by potentially masking the true outbreak size and duration through missing nodes and edges [60, 61]. Crucially, our evaluation framework treats the sampled empirical network as an absolute baseline. Consequently, while a generator like the DCSBM may closely fit the observed empirical structures, those underlying structures may themselves be incomplete or biased representations of the total population of interest.

Furthermore, because our five baseline networks were collected using identical geo-located proximity sensors over windows of several days using similar data collection protocols, it remains untested whether these generator rankings generalize to broader telemetry, observational, or self-reported data styles. Short observation windows may yield reasonably reliable degree-distributions, but accurately estimating community boundaries requires capturing enough repeated within-block contacts, just as calculating local clustering depends on capturing explicit triadic closures. The relative rankings among generators may therefore reflect a confluence of how well each generator’s signature features can be inferred from the available data alongside how those features are actually expressed in the true population. A further constraint is that the SEIR analysis relied on a single infectious period (*γ* = 2 days, the empirically established symptomatic-to-removal window for rabies in this setting), meaning that continuous *γ*-sensitivity probed by the SIS sweep is not directly mirrored in the SEIR comparison. Moreover, the Poisson-modeled latent period in the SEIR model (mean 30 days, standard deviation *≈* 5.5) captures most but not all of the reported variation, omitting the long tail of incubation periods extending up to 125 days [52]; however, this tail is unlikely to alter the comparative structural performance of the generators evaluated in this study. Lastly, the empirical canine rabies outbreaks are still simulation-based only and are not cross-referenced or correspond to real-life outbreaks, and may thus not necessarily be fully representative of observed canine rabies outbreak dynamics.

Beyond the substantive findings on generator choice, the data-driven kernel-density-valley approach used here to classify outbreaks as minor or major (see Materials and methods) is itself a small methodological contribution. The standard alternatives in the network-epidemic literature are arbitrary thresholds (for example, declaring any outbreak that infects more than 10% of the population a ‘major’ outbreak). The kernel-density approach instead locates the threshold at the natural valley between the bimodal density modes that arise in stochastic outbreak ensembles, which removes a researcher degree of freedom and is robust to the bandwidth choice across the range we tested. The approach should transfer to any network-epidemic study where the bimodal minor/major structure is present, and provides a more principled basis for outbreak classification in cross-generator comparisons of the kind performed here.

Several extensions follow naturally from this work. The most pressing is scaling these models up to city-level populations, the scale at which rabies vaccination programs typically operate but where generating networks with realistic edge and community densities becomes computationally demanding. A second concerns ecological fidelity: our unweighted proximity-network framework cannot capture the full social or spatial complexity of free-roaming dog systems, nor does it currently extend to companion-dog populations whose contacts are mediated by households, dog parks, and veterinary facilities. Extending this comparative framework to weighted, temporal, multilayer or scalable networks would represent a major step forward.

Finally, applying the framework to other proximity-transmitted infectious diseases, such as toxoplasmosis, brucellosis or Ebola, would clarify how broadly the generator rankings transfer to other disease systems.

Taken together, our results contribute three insights to the network epidemiology of contact-transmitted infectious diseases. First, they identify the degree-corrected stochastic block model as the most structurally and epidemiologically consistent of the five generators tested for reconstructing free-roaming dog contact networks, with its performance advantage arising from the joint representation of degree heterogeneity and community structure. Second, they provide an empirically grounded criterion — pathogen-dependent topological sensitivity — for determining when generator selection is consequential and when simpler generators suffice: for short-infectious-window pathogens such as canine rabies, generator choice can substantially shift cumulative incidence and outbreak-duration predictions, whereas for longer infectious windows, predictions gradually converge across generators to within roughly two percentage points of the empirical reference for a window of nine days. This indicates that generator choice matters most precisely for acute, rabies-like pathogens, while differences largely attenuate for longer-burning infections. Third, the results motivate a working hypothesis concerning the non-orthogonality of structural axes in network reconstruction: the implicit clustering that emerges from jointly modeling degree and community structure reproduces empirical network structure more faithfully than explicit clustering targeted in isolation, suggesting that frameworks that treat structural dimensions as independent may materially understate the difficulty of network reconstruction. Together, these findings support the use of degree-corrected stochastic block models for network-based modeling and intervention design in canine rabies and comparable infectious disease systems.

## Supporting information

**S1 Fig. Network topology summaries per graph type for different seed run sizes.**

**S2 Fig. Cumulative average of mean betweenness and degree.** Running means for all graph types for four seed sizes ranging between 100 and 5000.

**S3 Fig. Kolmogorov–Smirnov distances to the empirical degree distribution.** Distributions of the Kolmogorov-Smirnov distances between each synthetic degree distribution and the empirical degree distribution computed for all distributions of the different seed totals.

**S4 Fig. Schematic representations of the SIS (top) and SEIR (bottom) compartmental disease transmission models.** State variables and parameter values are defined in S2 Table and S3 Table.

**S5 Fig. Factorial outbreak sampling results.** Outbreak simulation outcomes under the factorial sampling design for the SIS and SEIR models at the *γ* values indicated in each panel. Panels show (from left to right) average prevalence, peak prevalence, and cumulative incidence for each of the five graph types in comparison to the empirical network. Only outbreaks with at least one secondary infection are shown.

**S6 Fig. Factorial sampling SEIR outbreak results.** SEIR final outbreak size and outbreak duration for three values of *β*, representing its full modeled range, separated into minor and major outbreaks for outbreaks simulated with each graph type and the empirical network. The x-axis shows outbreak duration measured in days (maximum = 1000) and the y-axis shows the final outbreak size measured as the proportion of removed individuals at the final day of the outbreak. Major outbreaks are indicated with darker shading compared to minor outbreaks.

**S7 Fig. Factorial sampling SIS outbreak results.** SIS model summary of prevalence and total infections for remaining gamma values highlighting the increase in prevalence and decrease in total number of cases as *γ* increases.

**S8 Fig. Summary of betweenness distributions across 5000 iterations of each graph type.** Each point on the x-axis summarizes the distribution of frequency values observed across all graphs of a given graph type for a specific betweenness value. The shaded band indicates the interquartile range (25th to 75th percentile) and the solid line shows the median.

**S9 Fig. SIS model summary of prevalence and total infections for remaining gamma values.**

These results highlight the increase in prevalence and decrease in total number of cases as *γ* increases.

**S10 Fig. Final outbreak size and outbreak duration of minor and major outbreaks for different transmission rates and for each graph type.**

**S11 Fig. Single graph representations of additional empirical contact networks and corresponding synthetic networks from each graph type.** Networks shown for: A) Habi, B) Hepang, C) Sabaneta, D) Romana.

**S12 Fig. Distributions of six network parameters for all additional locations.** The six measures include network density, global clustering, average betweenness, average degree, median degree, and maximum degree. Results are shown for all 5000 graph iterations of each graph type as well as their observed values in the empirical network for all four additional network locations.

**S13 Fig. Degree distribution over 5000 iterations of each graph type for all four additional network locations on a log-scale.**

**S14 Fig. Kolmogorov-Smirnov distances for the degree distributions of all additional locations.** The graph shows the distributions of the Kolmogorov-Smirnov distances between each synthetic degree distribution and the empirical degree distribution computed for all four additional network locations.

**S1 Table. Convergence analysis across graph types.** Standard error, stepwise mean drift and precision gain for mean degree and mean betweenness across sample sizes.

**S2 Table. State variables for the SIS and SEIR models. S3 Table. Parameter values for the SIS and SEIR models.**

**S4 Table. Definitions and epidemiological significance of calculated network metrics.**

**S5 Table. Size and duration of minor and major SEIR outbreaks.** Mean and standard deviation of final outbreak size and duration per graph type and transmission rate.

## Acknowledgments

The simulations presented in this manuscript were performed on the sciCORE scientific computing cluster of the University of Basel (http://scicore.unibas.ch/). We thank the data collection teams whose proximity-sensor fieldwork in Chad, Indonesia, and Guatemala made this study possible. We thank Dr. Mirjam Laager for the consultations on her published work and Professor Jakob Zinsstag for data access and consultation.

Several co-authors used AI assistance (Claude, Anthropic) during manuscript preparation. The AI helped with editorial review, prose refinement, citation research, figure-caption drafting, preparation of an implementation brief for the Newman clustered random graph generator that the lead author then coded in R, and identification of candidate peer reviewers. The authors reviewed all AI-assisted content. The AI system did not run simulations, generate empirical results, or access the contact-network data. All scientific claims and editorial decisions are the authors’.

## References

1. Craft ME. Infectious disease transmission and contact networks in wildlife and livestock. Philosophical Transactions of the Royal Society B: Biological Sciences. 2015 May;370(1669):20140107. Available from: https://royalsocietypublishing.org/doi/10.1098/rstb.2014.0107. doi:10.1098/rstb.2014.0107.

2. Keeling MJ, Eames KTD. Networks and epidemic models. Journal of The Royal Society Interface. 2005 Sep;2(4):295–307. Available from: https://royalsocietypublishing.org/doi/10.1098/rsif.2005.0051. doi:10.1098/rsif.2005.0051.

3. Laager M, Mbilo C, Madaye EA, Naminou A, Lèchenne M, Tschopp A, et al. The importance of dog population contact network structures in rabies transmission. PLOS Neglected Tropical Diseases. 2018 Aug;12(8):e0006680. Available from: https://dx.plos.org/10.1371/journal.pntd.0006680. doi:10.1371/journal.pntd.0006680.

4. Anderson RM, May RM. Infectious diseases of humans: dynamics and control. Reprinted ed. Oxford: Oxford Univ. Press; 2010.

5. Raynor B, De La Puente-León M, Johnson A, Díaz EW, Levy MZ, Recuenco SE, et al. Movement patterns of free-roaming dogs on heterogeneous urban landscapes: Implications for rabies control. Preventive Veterinary Medicine. 2020 May;178:104978. Available from: https://linkinghub.elsevier.com/retrieve/pii/S0167587719307196. doi:10.1016/j.prevetmed.2020.104978.

6. Sah P, Otterstatter M, Leu ST, Leviyang S, Bansal S. Revealing mechanisms of infectious disease spread through empirical contact networks. PLOS Computational Biology. 2021 Dec;17(12):e1009604. Available from: https://dx.plos.org/10.1371/journal.pcbi.1009604. doi:10.1371/journal.pcbi.1009604.

7. Shirley MDF, Rushton SP. The impacts of network topology on disease spread. Ecological Complexity. 2005 Sep;2(3):287–99. Available from: https://linkinghub.elsevier.com/retrieve/pii/S1476945X05000280. doi:10.1016/j.ecocom.2005.04.005.

8. Wilson-Aggarwal JK, Goodwin CED, Lèchenne M, Swan GJF, Sidouin MK, Silk MJ, et al. Spatial-temporal dynamics of contact among free-ranging domestic dogs Canis familiaris in rural Africa. bioRxiv. 2024. Available from: https://www.biorxiv.org/content/early/2024/06/29/2024.06.26.600798. doi:10.1101/2024.06.26.600798.

9. Watts DJ, Strogatz SH. Collective dynamics of ‘small-world’ networks. Nature. 1998 Jun;393(6684):440-2. Available from: https://www.nature.com/articles/30918. doi:10.1038/30918.

10. White LA, Forester JD, Craft ME. Using contact networks to explore mechanisms of parasite transmission in wildlife. Biological Reviews. 2017 Feb;92(1):389–409. Available from: https://onlinelibrary.wiley.com/doi/10.1111/brv.12236. doi:10.1111/brv.12236.

11. Bansal S, Grenfell BT, Meyers LA. When individual behaviour matters: homogeneous and network models in epidemiology. Journal of The Royal Society Interface. 2007 Oct;4(16):879–91. Available from: https://royalsocietypublishing.org/doi/10.1098/rsif.2007.1100. doi:10.1098/rsif.2007.1100.

12. Hethcote HW, Yorke JA. Gonorrhea Transmission Dynamics and Control. 1st ed. No. 56 in Lecture Notes in Biomathematics. Berlin, Heidelberg: Springer Berlin Heidelberg; 1984. doi:10.1007/978-3-662-07544-9.

13. Kojaku S, Hèbert-Dufresne L, Mones E, Lehmann S, Ahn YY. The effectiveness of backward contact tracing in networks. Nature Physics. 2021 May;17(5):652–8. Available from: https://www.nature.com/articles/s41567-021-01187-2. doi:10.1038/s41567-021-01187-2.

14. Pastor-Satorras R, Vespignani A. Epidemic Spreading in Scale-Free Networks. Physical Review Letters. 2001 Apr;86(14):3200–3. Available from: https://link.aps.org/doi/10.1103/PhysRevLett.86.3200. doi:10.1103/PhysRevLett.86.3200.

15. Halász V, Rocklöv J. Designing network based intervention strategies for epidemics of infectious diseases from edge based infection probability. Scientific Reports. 2026 Jan;16(1):808. Available from: https://www.nature.com/articles/s41598-025-33300-3. doi:10.1038/s41598-025-33300-3.

16. Salathè M, Jones JH. Dynamics and Control of Diseases in Networks with Community Structure. PLoS Computational Biology. 2010 Apr;6(4):e1000736. Available from: https://dx.plos.org/10.1371/journal.pcbi.1000736. doi:10.1371/journal.pcbi.1000736.

17. Weber N, Carter SP, Dall SRX, Delahay RJ, McDonald JL, Bearhop S, et al. Badger social networks correlate with tuberculosis infection. Current Biology. 2013 Oct;23(20):R915–6. Available from: https://linkinghub.elsevier.com/retrieve/pii/S0960982213011238. doi:10.1016/j.cub.2013.09.011.

18. Zagmutt FJ, Schoenbaum MA, Hill AE. The Impact of Population, Contact, and Spatial Heterogeneity on Epidemic Model Predictions. Risk Analysis. 2016 May;36(5):939–53. Available from: https://onlinelibrary.wiley.com/doi/10.1111/risa.12482. doi:10.1111/risa.12482.

19. Miller JC. Spread of infectious disease through clustered populations. Journal of The Royal Society Interface. 2009 Dec;6(41):1121–34. Available from: https://royalsocietypublishing.org/doi/10.1098/rsif.2008.0524. doi:10.1098/rsif.2008.0524.

20. Smieszek T, Fiebig L, Scholz RW. Models of epidemics: when contact repetition and clustering should be included. Theoretical Biology and Medical Modelling. 2009 Dec;6(1):11. Available from: https://tbiomed.biomedcentral.com/articles/10.1186/1742-4682-6-11. doi:10.1186/1742-4682-6-11.

21. Volz EM, Miller JC, Galvani A, Ancel Meyers L. Effects of Heterogeneous and Clustered Contact Patterns on Infectious Disease Dynamics. PLoS Computational Biology. 2011 Jun;7(6):e1002042. Available from: https://dx.plos.org/10.1371/journal.pcbi.1002042. doi:10.1371/journal.pcbi.1002042.

22. Farthing TS, Dawson DE, Sanderson MW, Lanzas C. Accounting for space and uncertainty in real-time location system-derived contact networks. Ecology and Evolution. 2020 Jun;10(11):4702–15. Available from: https://onlinelibrary.wiley.com/doi/10.1002/ece3.6225. doi:10.1002/ece3.6225.

23. Wilber MQ, Pepin KM, Campa H, Hygnstrom SE, Lavelle MJ, Xifara T, et al. Modelling multi-species and multi-mode contact networks: Implications for persistence of bovine tuberculosis at the wildlife–livestock interface. Journal of Applied Ecology. 2019 Jun;56(6):1471–81. Available from: https://besjournals.onlinelibrary.wiley.com/doi/10.1111/1365-2664.13370. doi:10.1111/1365-2664.13370.

24. Hampson K, Coudeville L, Lembo T, Sambo M, Kieffer A, Attlan M, et al. Estimating the Global Burden of Endemic Canine Rabies. PLOS Neglected Tropical Diseases. 2015 Apr;9(4):e0003709. Available from: https://dx.plos.org/10.1371/journal.pntd.0003709. doi:10.1371/journal.pntd.0003709.

25. Zinsstag J, Lechenne M, Laager M, Mindekem R, Naïssengar S, Oussiguèrè A, et al. Vaccination of dogs in an African city interrupts rabies transmission and reduces human exposure. Science Translational Medicine. 2017 Dec;9(421):eaaf6984. Available from: https://www.science.org/doi/10.1126/scitranslmed.aaf6984. doi:10.1126/scitranslmed.aaf6984.

26. Brookes VJ, Dürr S, Ward MP. Rabies-induced behavioural changes are key to rabies persistence in dog populations: Investigation using a network-based model. PLOS Neglected Tropical Diseases. 2019 Sep;13(9):e0007739. Available from: https://dx.plos.org/10.1371/journal.pntd.0007739. doi:10.1371/journal.pntd.0007739.

27. Guimerà R, Sales-Pardo M. Missing and spurious interactions and the reconstruction of complex networks. Proceedings of the National Academy of Sciences. 2009 Dec;106(52):22073–8. Available from: https://pnas.org/doi/full/10.1073/pnas.0908366106. doi:10.1073/pnas.0908366106.

28. Silk MJ, Jackson AL, Croft DP, Colhoun K, Bearhop S. The consequences of unidentifiable individuals for the analysis of an animal social network. Animal Behaviour. 2015 Jun;104:1–11. Available from: https://linkinghub.elsevier.com/retrieve/pii/S0003347215001025. doi:10.1016/j.anbehav.2015.03.005.

29. Bazzi M, Jeub LGS, Arenas A, Howison SD, Porter MA. A framework for the construction of generative models for mesoscale structure in multilayer networks. Physical Review Research. 2020 Apr;2(2):023100. Available from: https://link.aps.org/doi/10.1103/PhysRevResearch.2.023100. doi:10.1103/PhysRevResearch.2.023100.

30. Brask JB, Silk M, Weiss MN. An introduction to generative network models and their use in animal sociality research. Animal Behaviour. 2025 Dec;230:123364. Available from: https://linkinghub.elsevier.com/retrieve/pii/S000334722500291X. doi:10.1016/j.anbehav.2025.123364.

31. Staudt CL, Hamann M, Gutfraind A, Safro I, Meyerhenke H. Generating realistic scaled complex networks. Applied Network Science. 2017 Dec;2(1):36. Available from: https://appliednetsci.springeropen.com/articles/10.1007/s41109-017-0054-z. doi:10.1007/s41109-017-0054-z.

32. Vaca-Ramírez F, Peixoto TP. Systematic assessment of the quality of fit of the stochastic block model for empirical networks. Physical Review E. 2022 May;105(5):054311. Available from: https://link.aps.org/doi/10.1103/PhysRevE.105.054311. doi:10.1103/PhysRevE.105.054311.

33. Vu-Le TA, Anne L, Chacko G, Warnow T. EC-SBM synthetic network generator. Applied Network Science. 2025 May;10(1):15. Available from: https://appliednetsci.springeropen.com/articles/10.1007/s41109-025-00701-2. doi:10.1007/s41109-025-00701-2.

34. Erdős P, Rènyi A. On random graphs. Publicationes Mathematicae. 1959;6:290–7.

35. Barabási AL, Albert R. Emergence of scaling in random networks. Science. 1999 Oct;286(5439):509–12. doi:10.1126/science.286.5439.509.

36. Chung F, Lu L. The average distances in random graphs with given expected degrees. Proceedings of the National Academy of Sciences. 2002 Dec;99(25):15879–82. Available from: https://pnas.org/doi/full/10.1073/pnas.252631999. doi:10.1073/pnas.252631999.

37. Holland PW, Laskey KB, Leinhardt S. Stochastic blockmodels: First steps. Social Networks. 1983 Jun;5(2):109–37. Available from: https://linkinghub.elsevier.com/retrieve/pii/0378873383900217. doi:10.1016/0378-8733(83)90021-7.

38. Newman MEJ. Properties of highly clustered networks. Physical Review E. 2003 Aug;68(2):026121. Available from: https://link.aps.org/doi/10.1103/PhysRevE.68.026121. doi:10.1103/PhysRevE.68.026121.

39. Newman MEJ. Random Graphs with Clustering. Physical Review Letters. 2009 Jul;103(5):058701. Available from: https://link.aps.org/doi/10.1103/PhysRevLett.103.058701. doi:10.1103/PhysRevLett.103.058701.

40. Karrer B, Newman MEJ. Stochastic blockmodels and community structure in networks. Physical Review E. 2011 Jan;83(1):016107. Available from: https://link.aps.org/doi/10.1103/PhysRevE.83.016107. doi:10.1103/PhysRevE.83.016107.

41. Warembourg C, Fourniè G, Abakar MF, Alvarez D, Berger-González M, Odoch T, et al. Predictors of free-roaming domestic dogs’ contact network centrality and their relevance for rabies control. Scientific Reports. 2021 Jun;11(1):12898. Available from: https://www.nature.com/articles/s41598-021-92308-7. doi:10.1038/s41598-021-92308-7.

42. Lee C, Wilkinson DJ. A review of stochastic block models and extensions for graph clustering. Applied Network Science. 2019 Dec;4(1):122. Available from: https://appliednetsci.springeropen.com/articles/10.1007/s41109-019-0232-2. doi:10.1007/s41109-019-0232-2.

43. Wang YXR, Bickel PJ. Likelihood-based model selection for stochastic block models. The Annals of Statistics. 2017 Apr;45(2). Available from: https://projecteuclid.org/journals/annals-of-statistics/volume-45/issue-2/Likelihood-based-model-selection-for-stochastic-block-models/10.1214/16-AOS1457.full. doi:10.1214/16-AOS1457.

44. Lei J, Rinaldo A. Consistency of Spectral Clustering in Stochastic Block Models. The Annals of Statistics. 2015;43(1):215–37.

45. Qin T, Rohe K. Regularized Spectral Clustering under the Degree-Corrected Stochastic Blockmodel. In: Burges CJ, Bottou L, Welling M, Ghahramani Z, Weinberger K, editors. Advances in Neural Information Processing Systems. vol. 26. Curran Associates, Inc.; 2013. Available from: https://proceedings.neurips.cc/paper_files/paper/2013/file/0ed9422357395a0d4879191c66f4faa2-Paper.pdf.

46. Muse AH, Mwalili S, Ngesa O, Almalki SJ, Abd-Elmougod GA. Bayesian and Classical Inference for the Generalized Log-Logistic Distribution with Applications to Survival Data. Computational Intelligence and Neuroscience. 2021 Jan;2021(1):5820435. Available from: https://onlinelibrary.wiley.com/doi/10.1155/2021/5820435. doi:10.1155/2021/5820435.

47. Diekmann O, Heesterbeek H, Britton T. Mathematical Tools for Understanding Infectious Disease Dynamics. 1st ed. Princeton University Press; 2012. Available from: https://academic.oup.com/princeton-scholarship-online/book/13591. doi:10.23943/princeton/9780691155395.001.0001.

48. Singh R, Singh KP, Cherian S, Saminathan M, Kapoor S, Manjunatha Reddy GB, et al. Rabies – epidemiology, pathogenesis, public health concerns and advances in diagnosis and control: a comprehensive review. Veterinary Quarterly. 2017 Jan;37(1):212–51. Available from: https://www.tandfonline.com/doi/full/10.1080/01652176.2017.1343516. doi:10.1080/01652176.2017.1343516.

49. Müller T, Denzin N, Vos A, Freuling C. Incubation periods and mortality outcomes following rabies virus infection in mesocarnivorous reservoir hosts: implications for experimental design and veterinary policy – a review and meta-analysis. Virology Journal. 2025 Nov;22(1):374. Available from: https://virologyj.biomedcentral.com/articles/10.1186/s12985-025-02996-0. doi:10.1186/s12985-025-02996-0.

50. Tojinbara K, Sugiura K, Yamada A, Kakitani I, Kwan NCL, Sugiura K. Estimating the probability distribution of the incubation period for rabies using data from the 1948–1954 rabies epidemic in Tokyo. Preventive Veterinary Medicine. 2016 Jan;123:102–5. Available from: https://linkinghub.elsevier.com/retrieve/pii/S016758771530088X. doi:10.1016/j.prevetmed.2015.11.018.

51. Fekadu M, Shaddock JH, Baer GM. Excretion of Rabies Virus in the Saliva of Dogs. The Journal of Infectious Diseases. 1982;145(5):715–9.

52. Fekadu M. Canine Rabies. Onderstepoort Journal of Veterinary Research. 1993;60:421–7.

53. Lechenne M, Mindekem R, Madjadinan S, Oussiguèrè A, Moto DD, Naissengar K, et al. The Importance of a Participatory and Integrated One Health Approach for Rabies Control: The Case of N’Djamèna, Chad. Tropical Medicine and Infectious Disease. 2017 Aug;2(3):43. Available from: https://www.mdpi.com/2414-6366/2/3/43. doi:10.3390/tropicalmed2030043.

54. Mosleh M, Eckles D, Rand DG. Tendencies toward triadic closure: Field-experimental evidence. SocArXiv; 2024. Available from: https://osf.io/ys8zw. doi:10.31235/osf.io/ys8zw.

55. Broido AD, Clauset A. Scale-free networks are rare. Nature Communications. 2019 Mar;10(1):1017. Available from: https://www.nature.com/articles/s41467-019-08746-5. doi:10.1038/s41467-019-08746-5.

56. Davis S, Abbasi B, Shah S, Telfer S, Begon M. Spatial analyses of wildlife contact networks. Journal of The Royal Society Interface. 2015 Jan;12(102):20141004. Available from: https://royalsocietypublishing.org/doi/10.1098/rsif.2014.1004. doi:10.1098/rsif.2014.1004.

57. Hamede RK, Bashford J, McCallum H, Jones M. Contact networks in a wild Tasmanian devil ( *Sarcophilus harrisii* ) population: using social network analysis to reveal seasonal variability in social behaviour and its implications for transmission of devil facial tumour disease. Ecology Letters. 2009 Nov;12(11):1147–57. Available from: https://onlinelibrary.wiley.com/doi/10.1111/j.1461-0248.2009.01370.x. doi:10.1111/j.1461-0248.2009.01370.x.

58. Machens A, Gesualdo F, Rizzo C, Tozzi AE, Barrat A, Cattuto C. An infectious disease model on empirical networks of human contact: bridging the gap between dynamic network data and contact matrices. BMC Infectious Diseases. 2013 Dec;13(1):185. Available from: https://bmcinfectdis.biomedcentral.com/articles/10.1186/1471-2334-13-185. doi:10.1186/1471-2334-13-185.

59. Salathè M, Kazandjieva M, Lee JW, Levis P, Feldman MW, Jones JH. A high-resolution human contact network for infectious disease transmission. Proceedings of the National Academy of Sciences. 2010 Dec;107(51):22020–5. Available from: https://pnas.org/doi/full/10.1073/pnas.1009094108. doi:10.1073/pnas.1009094108.

60. Cross PC, Creech TG, Ebinger MR, Heisey DM, Irvine KM, Creel S. Wildlife contact analysis: emerging methods, questions, and challenges. Behavioral Ecology and Sociobiology. 2012 Oct;66(10):1437–47. Available from: http://link.springer.com/10.1007/s00265-012-1376-6. doi:10.1007/s00265-012-1376-6.

61. Gènois M, Vestergaard CL, Cattuto C, Barrat A. Compensating for population sampling in simulations of epidemic spread on temporal contact networks. Nature Communications. 2015 Nov;6(1):8860. Available from: https://www.nature.com/articles/ncomms9860. doi:10.1038/ncomms9860.

62. He P, Klarevas-Irby JA, Papageorgiou D, Christensen C, Strauss ED, Farine DR. A guide to sampling design for GPS-based studies of animal societies. Methods in Ecology and Evolution. 2023 Aug;14(8):1887–905. Available from: https://besjournals.onlinelibrary.wiley.com/doi/10.1111/2041-210X.13999. doi:10.1111/2041-210X.13999.

